# Unraveling the seasonal dynamics of ixodid ticks: A flexible matrix population model with delayed life history effects

**DOI:** 10.1101/2024.01.08.574636

**Authors:** Yngvild Vindenes, Atle Mysterud

## Abstract

Many vector-borne diseases are sensitive to changes in land use and climate, making it crucial to understand the factors that govern the vector populations. Ixodid ticks, which serve as vectors for multiple diseases, have a slow life cycle compared to many of their hosts. The duration of each active life stage (larvae, nymph, adult) varies greatly and depends on factors such as timing of questing and development, host availability throughout the seasons, and photoperiod-related behavioral and developmental diapause. Importantly, the observable questing population only represents a fraction of the total tick population and may include overlapping generations in each stage. Mathematical models are therefore essential to understand how complex life cycle transitions and host interactions impact the dynamics of the tick population. In this study, we present a flexible seasonal matrix model for ixodid ticks that feed on small and large hosts varying in seasonal availability. This model incorporates the delayed life history effects of overwintering and seasonal timing of feeding, density regulation through limited host capacity, and scramble competition among larvae and nymphs for small hosts. We extract the equilibrium seasonal numbers of questing, feeding, and emerging ticks for each life stage, as well as the seasonal patterns of host use. We also calculate key life history characteristics including the mean generation time, stable stage structure, and reproductive values. The baseline model represents a northern life history of the sheep tick (*Ixodes ricinus*) feeding on a seasonal small host and constant large host, which is compared to a scenario without host seasonality and a scenario representing a southern ecosystem. Our findings support the importance of small hosts in regulating tick populations, and highlight that feeding of larvae is a critical transition. Our analyses shed light on the complex mechanisms underlying the seasonal composition of the questing population, with its important implications for disease risk. The model can be applied to other ixodid tick species and provides a framework for future investigations into population dynamics under various tick and host scenarios.

## Introduction

Under global change, vector-borne diseases are becoming a growing threat to human and animal health (Jones et al., 2008; Ogden, 2017). Many of these diseases are transmitted by arthropod vectors, which carry pathogens from reservoir hosts to new hosts, potentially leading to pathogen spillover into human populations (Hill et al., 2005). Arthropod vectors vary in numerous aspects of their biology, including host specificity, degree of association with hosts, and host-seeking behavior (Ogden and Lindsay, 2016). Understanding the regulation of vector populations is crucial for predicting disease occurrences and implementing effective risk mitigation strategies. However, this task is challenging due to the diverse and complex life cycles of these vectors. While mosquito-borne diseases have received much research attention (Reiter, 2001), tick-borne diseases are not as well studied, despite the fact that ticks represent a distinct group of vectors with unique and varied ecological characteristics (Ogden and Lindsay, 2016). Argasid (soft) ticks, for example, predominantly dwell in close association with their hosts, often within nests, which provides some level of protection against environmental conditions. Conversely, ixodid (hard-bodied) ticks spend most of their life cycle off-host, in various stages including diapause, development, and questing (actively seeking hosts), leaving them more exposed to environmental factors (Kahl and Gray, 2023). To gain a comprehensive understanding of disease dynamics involving tick vectors, it is essential to develop population models that capture the seasonal dynamics of their multi-annual life cycles (Dobson et al., 2011).

In the northern hemisphere, pathogens causing Lyme disease, tick-borne encephalitis (TBE), anaplasmosis, and babesiosis are all transmitted by generalist species of ixodid ticks questing in the environment (Wu-Chuang et al., 2021). These species include *Ixodes ricinus* in Europe, West Asia, and North Africa, *I. persulcatus* in Asia and Eastern Europe, and *I. scapularis* and *I. pacificus* in North America (Franke et al., 2013). The life cycle of ixodid ticks consists of four main stages: egg, larva, nymph, and adult. To progress to the next stage or reproduce, the larva, nymph, and adult stages require a blood meal from a vertebrate host (Mannelli et al., 2012). Environmental changes can have different impacts on each life stage (Gatewood et al., 2009). Additionally, the synchrony in questing of tick larvae and nymphs can differ depending on local climate conditions (*I. ricinus*; Estrada-Peña and Estrada-Sánchez, 2014). Previous models for tick population dynamics have primarily focused on the influence of climate variables, such as temperature and saturation deficit, on questing and molting (*I. scapularis*: Ogden et al. (2005), *I. ricinus*: Dobson et al. (2011); Li et al. (2016)). While the significance of seasonality in tick questing and molting is well recognized, less is known about the role of seasonal changes in the main host species that regulate tick populations.

Several host species, particularly small mammals such as rodents and shrews, exhibit strong seasonality in abundance (Tkadlec and Zejda, 1998; Andreassen et al., 2021). These small mammals, along with ground-feeding birds, are the main hosts for tick larvae. Nymphs, on the other hand, feed on a wide range of hosts, including small mammals and birds, while adult female ticks require a blood meal from larger hosts, typically deer, to reproduce (Mannelli et al., 2012; Hofmeester et al., 2016; Estrada-Peña et al., 2016). Given that the population of larvae is typically an order of magnitude larger than that of nymphs (Randolph, 2004), an important regulatory factor for tick populations is the feeding success of questing larvae, which depends on the availability of small hosts. Most species of voles, mice, and shrews have a high reproductive capacity under favorable conditions and can increase by several orders of magnitude throughout the season (Andreassen et al., 2021). However, the reproductive season of small hosts starts relatively late in northern ecosystems (May/June), resulting in peak density occurring in the fall. Additionally, small mammal hosts are not accessible for tick feeding until they leave their nest or burrow, which typically takes around a month. Consequently, the lowest densities of small hosts coincide with the onset of larval questing in late spring. Thus, the peak availability of small mammal hosts may not necessarily align with the peak questing period of larvae or nymphs. Large hosts, such as deer, have lower reproductive capacities and therefore exhibit more stable population densities throughout the year. However, the availability of large hosts to ticks can also vary seasonally due to migration, as these hosts are highly mobile (Qviller et al., 2013). In northern ecosystems, birds display strong seasonality as many species are migratory.

Furthermore, the current models for tick populations do not consider the impact of overwintering status and time of feeding on tick survival, transition, and reproduction. These factors play a crucial role in diapause patterns and the duration of the tick life cycle (Gray et al., 2016; Grigoryeva and Shatrov, 2022). For instance, the population of questing nymphs can consist of individuals that have i) fed and molted earlier in the same year, ii) fed the previous year and overwintered as fed prior to molting, iii) fed and molted the previous year and overwintered as unfed, or a combination of these scenarios (Randolph et al., 2002; Grigoryeva and Shatrov, 2022). Tracking overwintering status is essential for understanding the dynamics because individuals that have overwintered may have different probabilities of survival, questing (if unfed), and molting (if fed) compared to those that hatched/molted/fed in the current year. For unfed ticks, for example, the stored fat reserves from the previous meal will be gradually depleted, resulting in fewer resources for overwintered individuals (Randolph et al., 2002; Randolph, 2004). These ticks may be more inclined to quest under adverse conditions compared to newly emerged ticks. Additionally, seasonal timing of feeding is important as photoperiod acts as a crucial cue for diapause (Belozerov et al., 2002; Gray et al., 2016). Ticks that feed before a certain threshold in the summer are more likely to initiate the molting process within the same year, whereas ticks that feed after this threshold are more likely to overwinter as fed and molt the following summer (Gray et al., 2016). Since the seasonal timing of feeding can have long-lasting effects on the probability of molting/reproducing, overwintering, and potentially other processes like survival, it is essential to incorporate it in a dynamical model.

We consider the aforementioned factors as crucial components of the life cycle of ixodid ticks and essential for understanding their regulation. To further investigate these dynamics, this study presents a seasonal matrix population model for ixodid ticks that incorporates the seasonal variation in host availability. Our model not only tracks the main life stages of ticks but also their status in terms of feeding, seasonal timing of feeding, and overwintering, including lasting effects of these events. This comprehensive approach allows for a detailed understanding of the mechanisms underlying the seasonal composition of tick populations. Density regulation is included through limited space on each available host, leading to host capacity being filled during the months with the highest questing numbers. To demonstrate the applicability and flexibility of our approach, we compare results for a baseline scenario of a northern population of *I. ricinus* feeding on a small seasonal host and a large non-seasonal host to a scenario with no seasonality in the small host, and to a southern scenario. This comparison highlights how the model can be adapted to different life histories and ecological conditions. Overall, our seasonal matrix population model provides a valuable tool for studying the dynamics of ixodid ticks, incorporating the effects of host availability and other key factors, and allowing for the exploration of various scenarios and species.

## Methods

### Life cycle of *I. ricinus*

Our aim is to provide a flexible matrix model that can be applied to generalist species of ixodid ticks, including *I. ricinus*, *I. persulcatus*, *I. scapularis*, and *I. pacificus* (Gray et al., 2016), that are primary vectors for *Borrelia burgdorferi* sensu lato, the causative agent of Lyme disease, and various other pathogens. Although these tick species share the same main life stages (egg, larvae, nymphs, and adults) and exhibit similar patterns of host selection throughout their ontogeny, they can show large differences in the duration and phenology of each life stage. For instance, *I. ricinus* can overwinter in all stages (Grigoryeva and Shatrov, 2022), whereas the eggs and fed adult stage of *I. persulcatus* cannot overwinter (Gray et al., 2016). Within each species, individual variation in stage transitions is primarily influenced by environmental factors and seasonal cues that affect the likelihood of questing in unfed stages and the molting process of fed stages (Randolph, 2004; Gray et al., 2016).

Our model is primarily based on the life cycle of *I. ricinus*, which shows large flexibility in stage transitions. In central and northern Europe, the molting process occurs during summer and fall, and takes several weeks to complete depending on temperature (Kahl and Gray, 2023). Ticks that feed in spring and early summer, prior to July, may proceed to molt in the same year and overwinter as unfed in the next stage (behavioral diapause) or potentially quest already the same fall and, if successful, overwinter as fed. However, ticks that feed later in the season typically postpone molting until the following summer, and enter developmental diapause after feeding (Randolph, 2004; Grigoryeva and Shatrov, 2022). Unfed individuals, particularly in the nymph and adult stage, can survive for several months before seeking their next blood meal (Randolph, 2004; Grigoryeva and Shatrov, 2022). They can enter behavioral diapause during winter or other periods of unfavorable questing conditions, such as heat spells in the summer. Some adult ticks can even survive for two winters without feeding, while unfed larvae and nymphs that have overwintered will die before the second winter unless they find a host (Grigoryeva and Shatrov, 2022). Due to the many potential overlaps of these processes behind transition probabilities between stages, the time required to complete the life cycle varies greatly depending on environmental conditions and location. The lifetime ranges from two to over six years, reflecting adaptations to live in environments of varying host availability and conditions. Furthermore, the plasticity of stage durations results in the potential for overlapping generations within the questing population each year (Bregnard et al., 2021).

The questing phenology of the active stages in *I. ricinus* also vary across different geographic regions. In northern Europe the main questing period for nymphs and adults occurs in the spring, while the peak activity of larvae is in summer (Dantas-Torres and Otranto, 2013). Conversely, in southern Europe, the main questing period for nymphs occurs much earlier during winter and early spring, with minimal overlap with the larvae (Dantas-Torres and Otranto, 2013). In northern and central Europe, a bimodal pattern is often observed in the questing activity of nymphs and adults, with a spring peak followed by a smaller peak in the fall (Wongnak et al., 2022). This pattern may arise from different mechanisms, or combination of mechanisms. For example, poor questing conditions during the summer may increase the likelihood of behavioral diapause in the questing population. Alternatively, the different peaks could reflect the emergence of different generations of ticks, with a spring peak arising from overwintered unfed nymphs and adults, and a fall peak consisting of ticks that fed the previous year and molted in the current year to emerge in late summer. Furthermore, the fall peak could also include individuals that fed early in the same year and completed the molting process in time to quest in the fall (Bregnard et al., 2021). Following a blood meal the ticks can either initiate the molting process to the next stage or enter developmental diapause, a period of developmental arrest lasting several months, helping them to survive winter (Gray et al., 2016; Grigoryeva and Shatrov, 2022).

## Model description

In this section we describe the model and its main components, as well as our choices for parameter values in the baseline scenario for *I. ricinus* ticks in a northern region. Importantly, the model can be adjusted to fit the wide range of variation in local tick life histories and host availability by modifying parameters. A description of key model parameters can be found in Table 1, and additional details are given in the Supplementary Information.

**Table 1:** Overview of main parameters in the model, and whether they are constant or seasonal in the baseline model. Tick stage *j* refers to the relevant stage out of the 17 stages described in fig. 1. Transitions to relevant ‘previous’ stages occur between December and January.

| Parameter | Description | Seasonal or constant |
| --- | --- | --- |
| $a_{ji}$ | Survival probability stage $j$, month $i$ | Seasonal |
| $q_{ji}$ | Questing probability stage $j$, month $i$ | Seasonal |
| $m_{ji}$ | molting probability stage $j$, month $i$ | Seasonal |
| $h_{ji}$ | Hatching probability stage $j$, month $i$ | Seasonal |
| $r_{ji}$ | Reproduction probability stage $j$, month $i$ | Seasonal |
| $S_i$ | Small host availability in month $i$ | Seasonal |
| $L_i$ | Large host availability in month $i$ | Constant |
| $p_j$ | Small host preference in tick stage $j$ | Constant |
| $f_j(S_i)$ | Probability of finding a small host in tick stage $j$ | Seasonal |
| $f_j(L)$ | Probability of finding a large host in tick stage $j$ | Constant |
| $MS_j$ | Maximum probability of finding small host, tick stage $j$ | Constant |
| $ML_j$ | Maximum probability of finding large host, tick stage $j$ | Constant |
| $C$ | Capacity (feeding spots) per small host | Constant |
| $C_j$ | Capacity (feeding spots) per large host for stage $j$ | Constant |

### (a) Stage structure and population dynamics

The tick model is female-based with a monthly time scale. The tick population is classified into 17 distinct stages (fig. 1, table S1), reflecting not only the main life stage (egg, larvae, nymph or adult) but also the feeding state (unfed or fed), seasonal timing of feeding (spring or fall), and overwintering state (current or previous). The model uses a post-reproductive census, where individuals in each stage are counted at the start of each month. In the next month individuals may survive and make a transition to a different stage or reproduce, depending on the month and their current stage. For instance, eggs may hatch, unfed individuals may quest and feed, fed larvae and nymphs may molt, and fed adults may reproduce. Reproducing adults lay their eggs at the start of the next month and then die right before the subsequent census. We assume that adults feeding in the current month cannot reproduce until at least one month has passed since they first entered the fed stage. This effectively ensures a minimum two-month interval between feeding and egg-laying. Additionally, the feeding process incorporates density dependence through the availability of small and large hosts, described in the next subsection.

**Figure 1.**
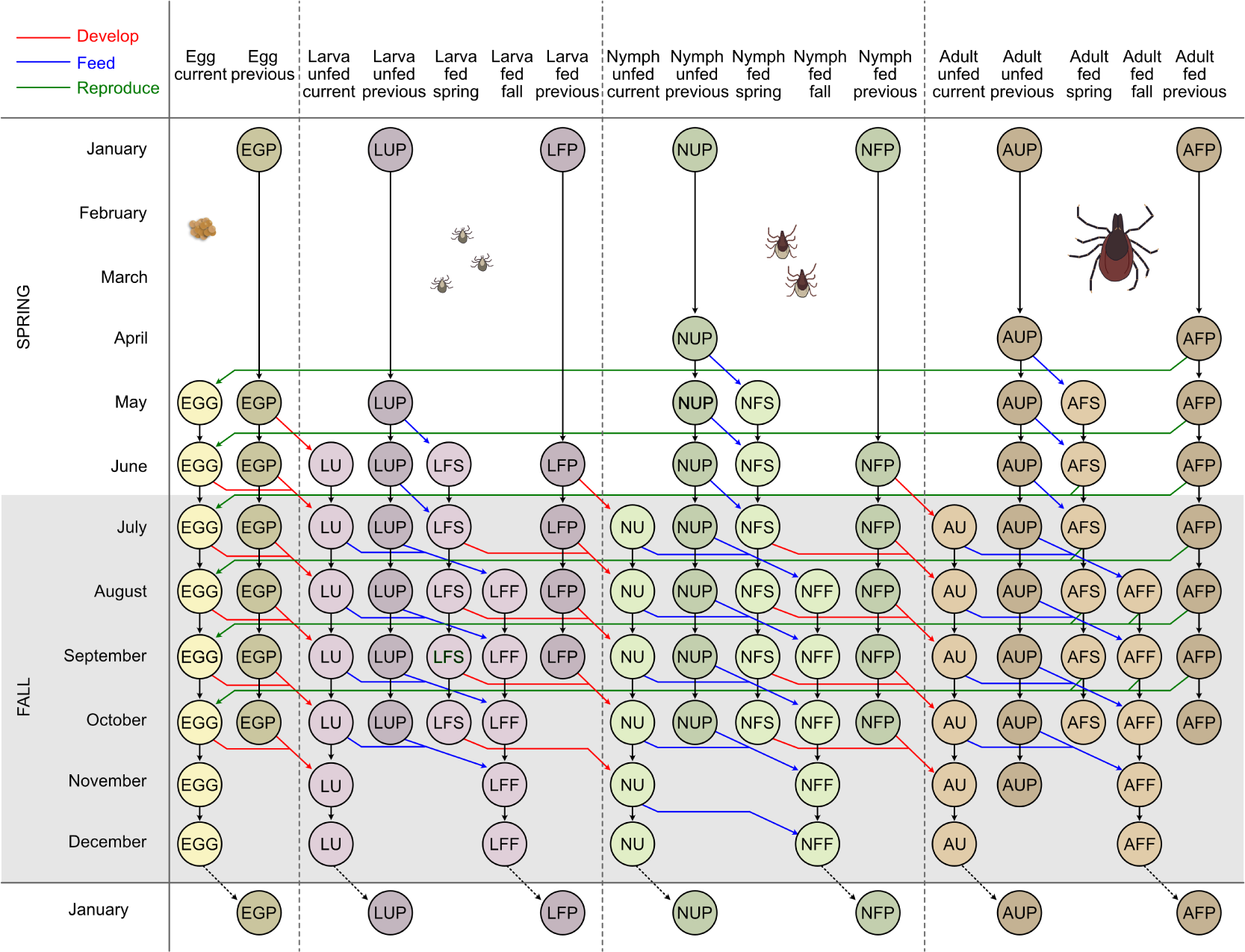
Overview of monthly transitions in the tick model representing the baseline northern scenario. Developmental transitions are shown in red, and represent hatching for eggs and molting for larvae and nymphs. Feeding transitions are shown in blue. The probability of feeding in each stage is density dependent and is determined by the questing probability and by the availability of small and large hosts. Reproduction is shown by green. Fed females that reproduce die after laying eggs. We separate between individuals that fed in spring or fall, determined by a threshold set to July in the baseline scenario. Fall fed individuals are more likely to enter diapause to overwinter and molt the next season. Similarly, individuals that emerge from molting in fall are more likely to overwinter as unfed and quest the following year, although some will quest in fall. Overwintered stages (both fed and unfed) are denoted as ‘previous’.

Denoting the stage-specific tick population vector at time step *t* and month *i* (*i* = 1, 2*, …,* 12) as **n***_i,t_*, the population size the next month is given by **n***_i_*_+1_*_,t_*_+1_ = **A_i_**,**_t_**(*S_i_, L_i_,* **n_i_**,**_t_**)**n_i_**,**_t_**, where **A_i_**,**_t_**(*S_i_, L_i_,* **n_i_**,**_t_**) is the projection matrix for month *i* at time *t*. This matrix depends on the availability of small hosts *S_i_*, the availability of large hosts *L_i_*, and on the current density of ticks **n_i_**,**_t_**, which together determine the transition probabilities from unfed to fed stages (described in next subsection). The monthly projection matrix can further be decomposed as **A_i_**,**_t_**(*S_i_, L_i_,* **n_i_**,**_t_**) = **T_i_**(*S_i_, L_i_,* **n_i_**,**_t_**)diag(**a***_i_*) + **F_i_**, where **F_i_** is a density independent fertility matrix for month *i*, describing reproduction from the three adult fed stages, **a***_i_* is the vector of stage-specific background survival probabilities, and **T_i_**(*S_i_, L_i_,* **n_i_**,**_t_**) is a density dependent transition matrix for month *i*.

### (b) Host availability and feeding of ticks

In the scenarios analyzed in this study, we consider that the small host availability *S_i_* shows seasonal variation, while the large hosts availability *L_i_* remains constant throughout the year (fig. 2). The transition of larvae, nymphs and adults from unfed to fed stages in a given month *i* depends on four main processes: i) questing, ii) host type preference, iii) the probability of finding a preferred host, and iv) available space to feed given that a host is found. The probability of questing in month *i* is denoted *q_ji_*, where the stage *j* represents one of the unfed stages (*LU*, *LUP*, *NU*, *NUP*, *AU*, *AUP*). The small host preference *p_j_* denotes the average proportion of the questing population within each stage that ‘search’ for small hosts, while the remaining proportion seeks large hosts (*p_L_* = 0.9, *p_N_* = 0.5, *p_A_* = 0 for larvae, nymphs, and adults, respectively). Importantly, this preference parameter does not imply that individual ticks actively search for a specific host type. Instead, it reflects the average outcomes of stage-specific questing behaviors, such as vertical positioning within the vegetation. For instance, larvae tend to quest at lower heights, where they are more likely to encounter small hosts (Dantas-Torres and Otranto, 2013). The stage-specific probability of finding a host of a given type is modeled as a logistic function of host availability, with the maximum probability depending on stage and host type (fig. 2b). The probability of a larvae finding a small host is denoted as *f_L_*(*S_i_*), the probability of a larva finding a large host is denoted as *f_L_*(*L_i_*), and similar notation is used for the other tick stages. We assume that larvae have a lower probability of finding a host or reaching a suitable spot on a host compared to nymphs and adults, due to their limited mobility.

**Figure 2.**
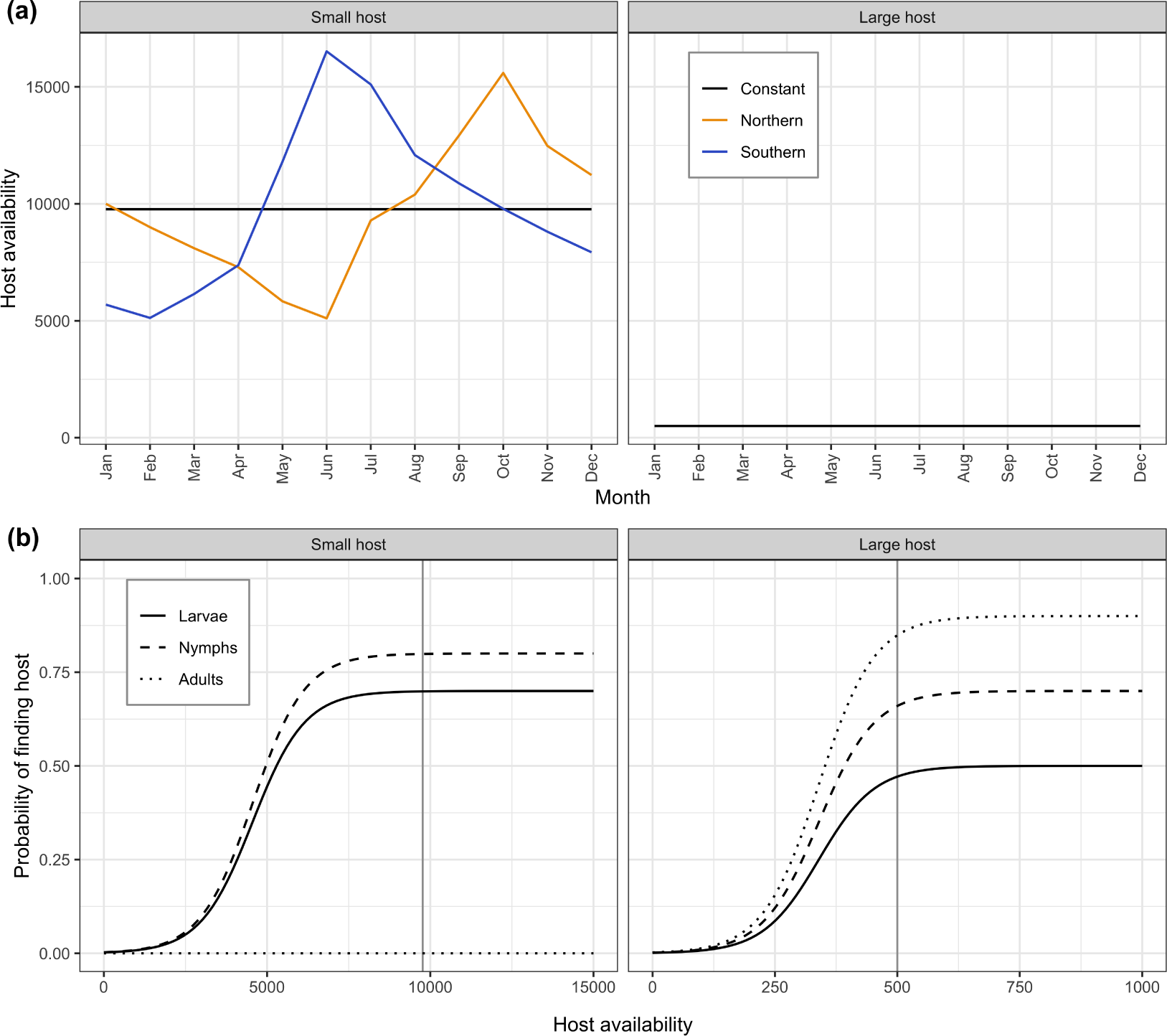
Properties of host availability and feeding used in the analyses. (a) Seasonal availability of small and large host availability used in the different scenarios. The small host availability represents adult individuals of a small mammal. (b) The stage-dependent probability of ticks finding a small host and large host used in the analyses. Vertical lines represent the mean host availability across scenarios.

Whether all ticks that find a host in a given month can feed depend on the host capacity. The total number of unfed larvae that find a small host in month *i* is 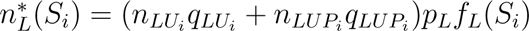, while the number of larvae finding a large host is 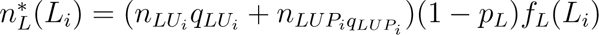 (similar expressions apply to the nymphs and adults). We assume that each small host has a capacity of *C* ‘feeding spots’. The total available spots on small hosts are distributed among the nymphs and larvae that find a small host, where nymphs are assumed to require three feeding spots each and larvae one spot each. The available small host spots are divided among the larvae and nymphs according to their relative proportions in the total requested spots. If the number of requested spots is below the total number of available spots (if 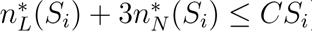) all ticks that find a host will feed. If not, the number of spots allocated to larvae is 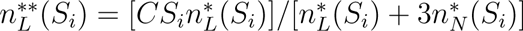 while the number allocated to nymphs is 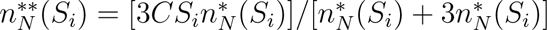. In our analyses we assume *C* = 500, meaning that each small host can feed 500 larvae or 167 nymphs per month, or a combination of both in between. This corresponds approximately to reported numbers on mice and voles scaled up to one month (Lindsø et al., 2023). As the population of questing larvae is typically much higher than that of questing nymphs, during months of high competition s, larvae will secure most of the available feeding spots on small hosts. For large hosts, the different tick stages are assumed to use different parts of the host (Mysterud et al., 2014) so that ticks only compete within stage. Each large host has a capacity *C_L_* for larvae, *C_N_* for nymphs, and *C_A_* for adults (all set to 5000 in our analyses).

### (c) Baseline northern scenario

The baseline parameter values for the model (fig. S5) are chosen to represent a northern population characterized by high survival rates due to low risk of desiccation, but slow development rates due to low mean temperature (Estrada-Peña and Estrada-Sánchez, 2014). The population is strongly influenced by seasonality, with most activity occurring from April to October. The peak questing occurs in May for nymphs, June for adults, and July for larvae, with smaller peaks in the fall representing questing from newly emerged individuals (fig. S5). Nymphs and adults are assumed to have slightly longer questing periods each year than larvae. Late summer is assumed to be the peak season for molting and egg laying by adult fed females (fig. S5). We assume each reproducing female lays 1500 eggs, in line with reported average values for *I. ricinus* (Gray, 1981; Grigoryeva and Shatrov, 2022).

The chosen baseline values for survival are based on information from Grigoryeva and Shatrov (2022) and Randolph (2004). We assume the survival is slightly higher in fed compared to unfed stages, as questing and feeding is associated with a higher risk of mortality. Survival estimates from empirical studies are often based on placing ticks in containers in the field, and may overestimate survival compared to natural populations due to the prevention of some mortality during feeding or predation (Randolph, 2004). We assume that overwintered stages have a progressively lower survival rate throughout the year as their stored resources are depleted. Survival in these stages eventually reaches zero, meaning overwintering individuals will die during the following year unless they successfully feed or reproduce in the case of adult fed females.

Photoperiod-induced diapause is accounted for through the separation of spring- and fall-fed ticks, which have different probabilities of molting. In January, fed ticks that have not molted will enter the ‘fed previous’ stage. July is set as the threshold for spring- and fall-feeding, corresponding to Northern Europe (Gray et al., 2016). Ticks that feed before July can molt later in the same year, then overwinter as unfed and quest next year. Larvae and nymphs that feed from July onwards are assumed to overwinter as fed, and complete the molting process in the following summer. These individuals may then quest in fall that year or overwinter as unfed, generating a one year delay in the life cycle (Grigoryeva and Shatrov, 2022). Adults that feed in spring are assumed to lay their eggs the same year, along with a few adults that feed in fall. But in the baseline scenario we assume most of the fall-fed adults will overwinter and lay their eggs next year.

### (d) Alternative southern scenario

As a comparison we created a scenario with parameters values corresponding to the southern range of the distribution of *I. ricinus* in Europe. Information on questing times was obtained from Dantas-Torres and Otranto (2013) who reported results from a forest location in southern Italy. The questing phenology is very different for this population compared to the north, with peak questing of nymphs in winter and early spring, of adults in early spring, and of larvae in summer (figs. S34, S35). Questing within each stage is also more spread out across months compared to the northern scenario (fig. S35). While information was not available on molting and survival for the Italian population, the authors suggest that the life cycle may be completed in about one year (Dantas-Torres and Otranto, 2013). High temperatures should allow fast development compared to the north, so we assume for this scenario that eggs that are laid in spring hatch to larvae that quest the same year. We further assume that the larvae can feed and develop to nymphs in the same year, with emergence in late fall and winter (figs. S34, S35). Nymphs quest in winter and early spring, molt to adults in the following summer and fall, then these adults then quest next spring. This leads to a life cycle closer to two years than one, but still very fast compared to the northern baseline scenario.

For the analyses in this scenario we also assume that the reproduction of the small host starts earlier in the season, and stops in summer when the temperature is high (Andreassen et al., 2021), so that the peak small host availability occurs earlier in the year (in June) compared to the baseline northern scenario (fig. 2a). The large host availability is assumed to be the same as in the northern scenario.

## Model outputs

The model is defined and analyzed in R (R Core Team, 2023). For each scenario and parameter combination we project the monthly population growth for several years, tracking each stage and also storing each monthly transition matrix, until well after equilibrium stable dynamics have been reached (visually assessed). When the equilibrium is reached, the monthly projection matrices are the same each year and the population will show only seasonal fluctuations (no interannual fluctuations). While it is theoretically possible to obtain multi-annual cycles with this model, most parameter combinations including all considered here will lead to a stable equilibrium with only seasonal cycles.

At equilibrium we extract the seasonal numbers of questing, feeding, fed and emerging ticks from each of the main active stages (larvae, nymphs, and adults). For the seasonal questing numbers we separate between different generations (current or previous). For the monthly numbers of successfully feeding individuals we separate between host type (small or large), and we also calculate the monthly degree of host use which indicates months of density regulation (Supplementary Information). Regarding the fed population, we separate individuals that fed in spring, fall, and the previous year. We also track the origin of newly emerging (molted) individuals each month, whether they come from overwintered individuals that fed the previous year, or from individuals that fed earlier in the same year.

For the equilibrium populations we calculate the annual projection matrix and its main components, and use these to calculate the stable stage structure **u**, reproductive values **v**, and generation time *G*. We also verify that the annual growth rate *λ* and the net reproductive rate *R*_0_ are equal to 1 (due to density regulation). For instance, the annual projection matrix from August to August is given by **A**_August-to-August_ = **A**_7_**A**_6_**A**_5_**A**_4_**A**_3_**A**_2_**A**_1_**A**_12_**A**_11_**A**_10_**A**_9_**A**_8_, where **A***_i_* is the extracted projection matrix for month *i*. The stable stage structure **u** and the reproductive values **v** are calculated as the right and left eigenvector of the projection matrix **A**_August-to-August_ associated with the dominant eigenvalue (Caswell, 2001). We do the same calculation for annual matrices with different starting month to obtain estimates for different months (Supporting Information). To extract the annual survival/transition matrix **U**_August-to-August_ we use the same approach as for the annual projection matrix (multiplying monthly survival/transition matrices extracted at equilibrium). The annual fertility matrix is **^F^**August-to-August ^=^ **^A^**August-to-August *^−^* **^U^**August-to-August ^(Caswell, 2009). Generation time is^ calculated as the mean age (in years) of reproducing females for the population at the stable equilibrium (Bienvenu and Legendre, 2015), inline6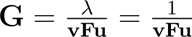. Since the females die after reproduction, the mean generation time also represents the mean lifetime in this case.

## Results

### (a) Baseline northern scenario

The baseline northern population has a generation time of 3.72 years at equilibrium. During the peak questing period, the number of questing larvae is an order of magnitude higher than the number of nymphs, which is in turn an order of magnitude higher than the number of adults (Fig. 3a). The peak month of the questing population mostly corresponds to the peak month of questing probability (Fig. S5), but there is a slight difference for larvae as their peak questing occurs in June while the peak questing probability is in July. The majority of questing ticks are unfed individuals that hatched or molted in the previous year, but there is also a smaller fall peak of questing individuals from the current year. Larvae mainly feed on small hosts as expected (fig. 3b), and most of their feeding occurs in July and August, after the peak questing period. Since the small host population is bigger in fall than spring (Fig. 2a), more questing larvae will successfully feed in fall. In June and July, when competition for small hosts is highest (figs. S12-S14), the probability of finding a small host is lower (fig. 2), resulting in a smaller proportion of questing larvae feeding in these months. July has the highest number of feeding larvae because a large number are questing relative to the available number of small hosts. In most months, the number of nymphs feeding on small hosts is higher than the number feeding on large hosts, except in June and July when competition with larvae is high. In the other months, all questing ticks that find a host will successfully feed. The adult ticks only feed on large hosts and their peak feeding period corresponds to the peak in questing (fig. 2b). The peaks for larvae and nymphs feeding on large hosts also align with the peak in questing, as there is no competition for these hosts in the baseline scenario. The stable population structure at equilibrium is skewed towards the egg and unfed larvae (fig. S8), while the reproductive value increases with each life stage and is highest for adults (fig. S9). Furthermore, adults that feed in fall have a higher reproductive value compared to adults that feed in spring (fig. S9).

**Figure 3.**
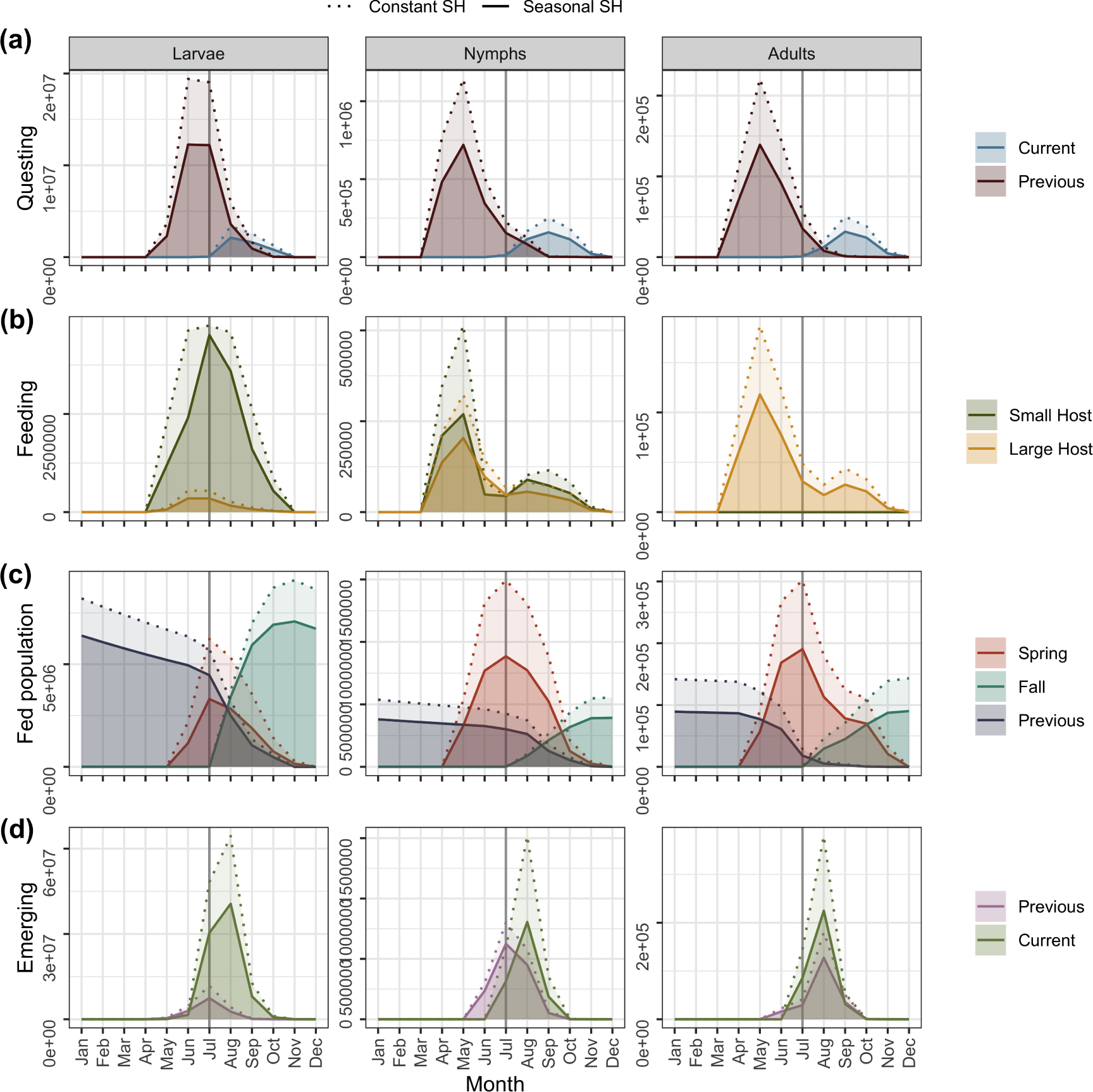
Model outputs at the equilibrium stable state in the baseline northern scenario with a seasonal small host (solid lines) and the corresponding outputs for the same model with a constant small host (dashed lines). (a) Questing population of larvae, nymphs and adults from individuals that emerged as unfed in the current year and from individuals that emerged in the previous year and overwintered as unfed (previous). (b) Numbers of larvae, nymphs and adults that feed on each host type each month. (c) Population of fed larvae, nymphs and adults divided into those that fed in spring (before July) the same year, fall (from July onwards) the same year, or previous year (previous). (d) Numbers of emerging (unfed) larvae, nymphs and adults each month, divided into those that fed earlier in the same year (current) and those that fed previous year and overwintered as fed (previous).

The population of fed individuals at equilibrium also varies throughout the year (fig. 3c), increasing as newly fed individuals enter and decreasing due to mortality and transitions out of the fed stages. The individuals that survive and feed in a given month are included in the fed population the following month, categorized as either spring-fed or fall-fed based on their seasonal timing of feeding. The spring-fed population reaches its peak during the summer, and these individuals undergo molting (or reproduction) in the summer or fall of the same year. All fall-fed larvae, nymphs, and most fall-fed adults enter diapause and complete their molting (or reproduction) in the following year. Figure 3d shows the number of newly emerging (unfed) larvae, nymphs, and adults each month, along with whether they originate from overwintered eggs or previously fed individuals (“Previous”), or from eggs laid and individuals fed in the current year (“Current”). The majority of the larvae emerge from eggs laid in the current year, while approximately half of the nymphs emerge from larvae that were fed earlier in the same year, and about half of the adults from nymphs that were fed earlier in the same year.

### (b) Varying host levels and preferences

The tick population is primarily limited by the availability of small hosts, rather than large hosts (fig. 4). Some large hosts are required for reproduction since adult ticks only feed on large hosts. However, increasing the availability of large hosts beyond a certain level has no impact on the tick population size (when the small host availability follows the baseline scenario, fig. 2), while changing the availability of small hosts directly affects the tick population (fig. 4).

**Figure 4.**
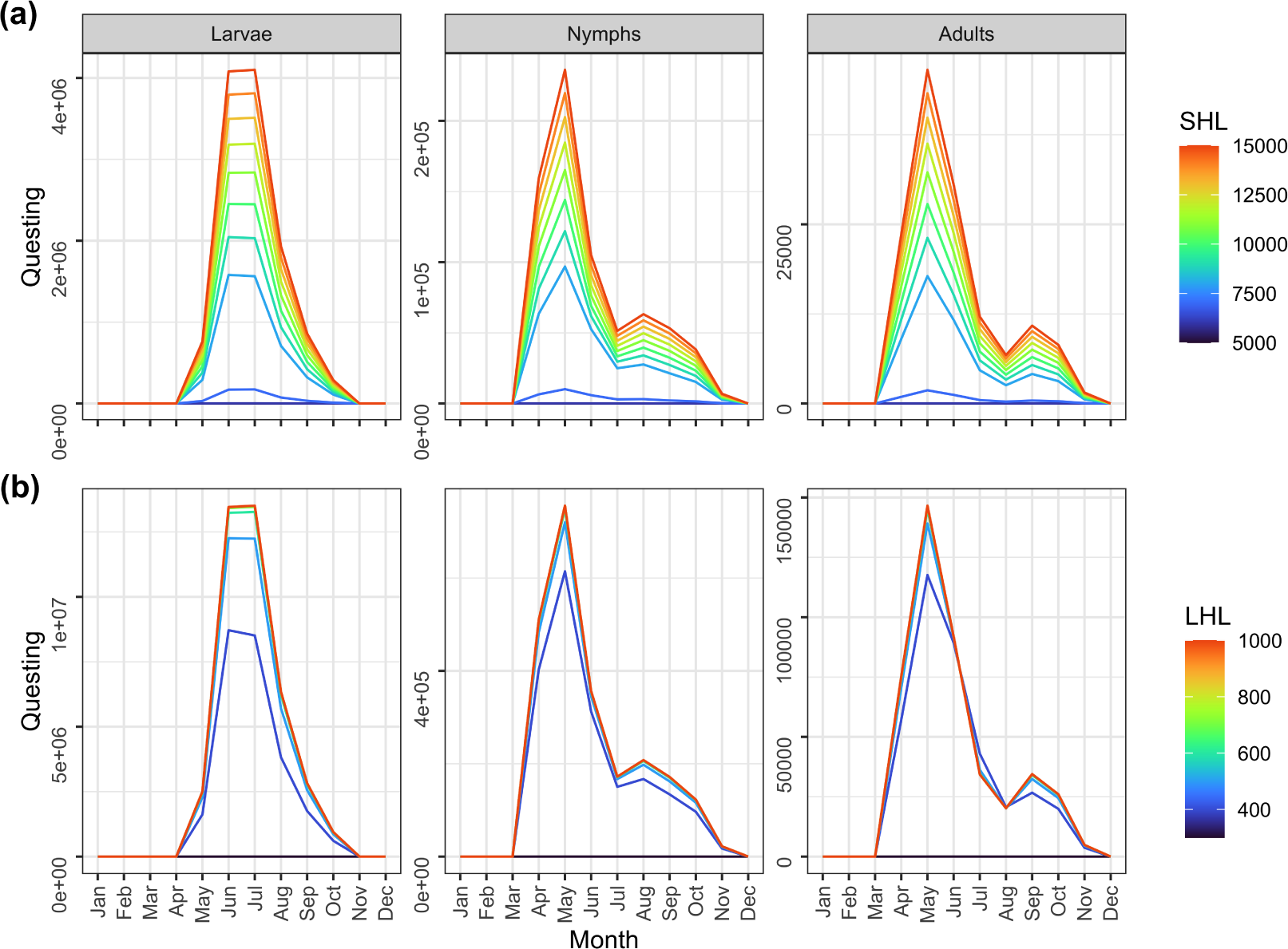
Effects of varying host availability on the equilibrium questing population in the baseline northern scenario. (a) Effects of varying small host level (SHL values refer to the initial January size). The large host availability is 500 as in the baseline scenario. (b) Effects of varying the large host level (LHL) which is constant across seasons. The small host availability is seasonal, as shown in figure 2.

The tick population is also more sensitive to the small host preference of larvae compared to nymphs (fig. 5). If the small host preference in larvae is very low (below approximately 0.5, which seems unrealistic), the tick population is not viable under the baseline scenario. As long as the tick population is viable, an intermediate value of the small host preference in larvae leads to the largest tick population size (fig. 5). This intermediate value allows for the optimal distribution of questing larvae among small and large hosts, resulting in a larger tick population. Additionally, the competition with nymphs for small hosts is reduced during the critical months June and July, further contributing to an increase in the tick population. On the other hand, changing the small host preference of nymphs has a smaller effect (fig. 5). Increasing nymphal preference leads to a higher equilibrium number of questing larvae and adults, but slightly fewer questing nymphs. As more nymphs feed on small hosts, the competition with larvae will increase during months when questing is high relative to host availability (given the baseline larval preference of 0.9), and since there are more questing larvae overall, they are more likely to obtain a larger share of the available spots on small hosts.

**Figure 5.**
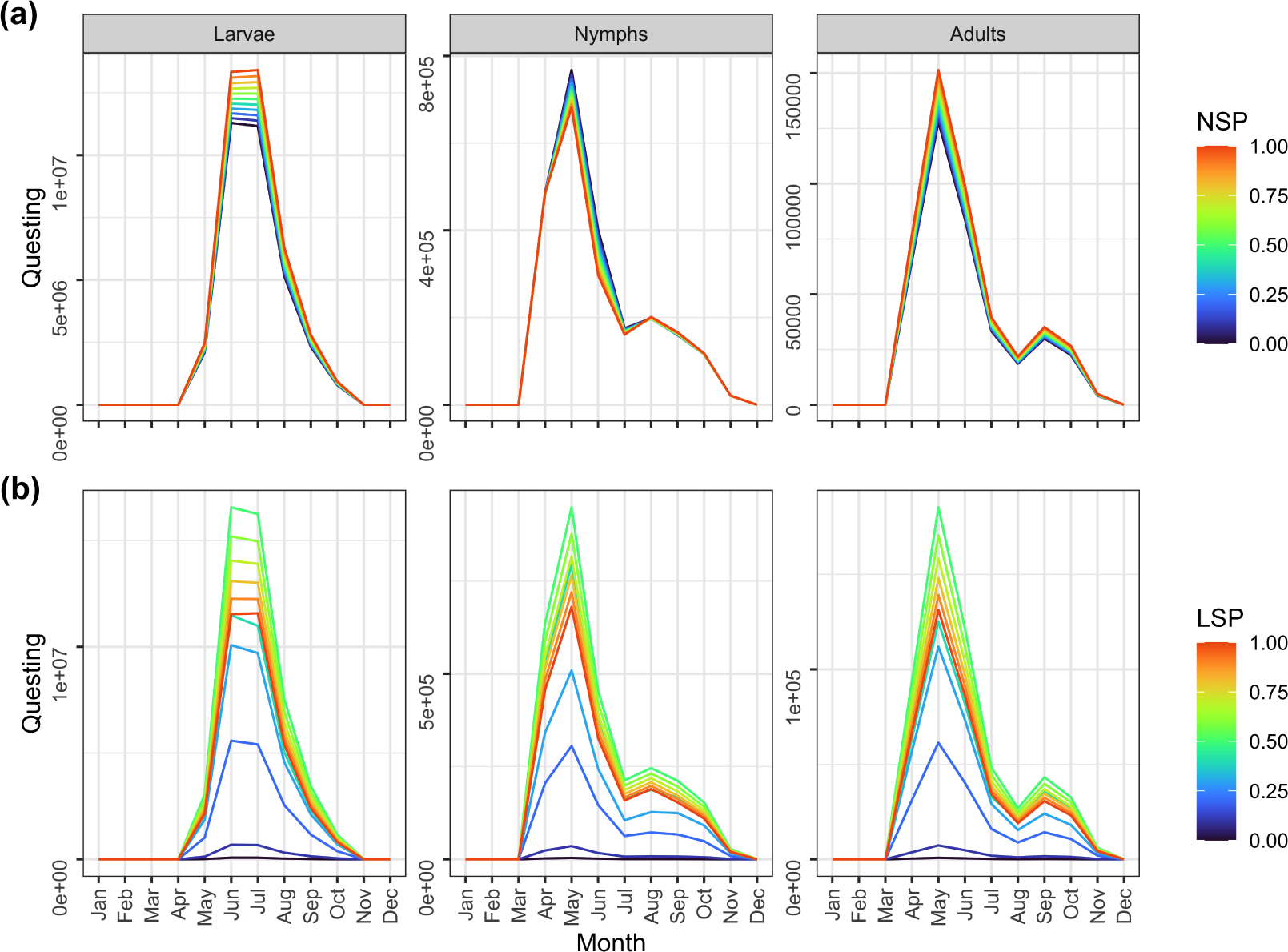
Effects of variation in small host preference of nymphs and larvae on the equilibrium questing population in the baseline northern scenario. Adults feed only on large hosts. (a) Effects of varying nymph small host preference (NSP). The larval small host preference is here 0.9. (b) Effects of varying larval small host preference (LSP). The nymph small host preference is here 0.5. Results corresponding to larval host preference lower than about 0.4 lead to declining, non-viable populations (no equilibrium reached, results shown are after 1000 time steps).

### (c) Constant small host

Comparing the baseline northern model to a model with constant small host (Supporting Information S5), the main effect is an increased equilibrium tick population due to more larvae and nymphs being able to feed during months of high questing activity. In addition, the small host capacity is completely filled for a longer time (June, July, and August), leading to increased competition between larvae and nymphs during these months.

### (d) Southern scenario

The results for the southern scenario are presented in the Supporting Information (Supporting Information S6). In this scenario, the mean generation time at equilibrium is 2.2 years, which is much shorter than the 3.7 years in the northern scenario. Furthermore, the peak availability of small hosts occurs earlier in the year (fig. 2a) in comparison to the northern scenario, enabling more larvae to find a host and feed during their peak questing season. Since nymphs and larvae quest at different times of the year, there is less competition among these stages for the small hosts. However, in the months of July, August, and September, all of the small host capacity is filled, indicating a strong competition among larvae (figs. S42-S44). Note that in this scenario, only larvae use their entire available host capacity during these critical months.

## Discussion

Ixodid ticks play a crucial role as vectors for various zoonotic pathogens that are emerging due to global change (Jones et al., 2008). However, many important stages of their life cycle remain hidden and cannot be directly observed in the field (Dobson, 2014). The visible questing population of these ticks represents only a small fraction, and traditional methods like cloth-lure and dragging techniques often underestimate their actual numbers (Nyrhilä et al., 2020). Understanding the underlying mechanisms behind seasonal numbers of questing ticks, with the close links to disease hazard (Ostfeld et al., 2006), is therefore an important modeling task (Dobson et al., 2011). In this study we have developed a seasonal matrix model specifically designed to capture the delayed effects of feeding and overwintering timing, while also accounting for the seasonality of host availability. This model is flexible and can be adjusted to accommodate the wide range of variation in life history and seasonality observed in different species of ixodid ticks and vertebrate hosts. We presented results from a northern (baseline) and a southern scenario, focusing on *I. ricinus* ticks in Europe. Our model sheds light on the often overlooked role of hosts and their seasonality in regulating tick populations, and also demonstrates how the multi-annual life cycle of ticks, with its unobservable stages, impacts the seasonal dynamics. Although gathering empirical estimates for all parameters is challenging for any tick dynamics model, our approach provides a valuable tool for exploring how different scenarios of life history can affect the questing population and its composition.

A key novel aspect of our model is its ability to track the state of individuals with respect to overwintering, feeding, and seasonal timing of feeding, and to incorporate lasting effects of these states on individual performance for the next months and year. This feature provides a more detailed understanding of the composition of the questing population throughout the season, not only in terms of the main life stages but also with respect to to different generations within each life stage, relevant for understanding the transmission of pathogens. While other models have included photoperiod-induced diapause (e.g., Dobson et al., 2011), to our knowledge this is the first model for ixodid ticks that includes lasting effects of overwintering and seasonal timing of feeding, affecting the performance of individuals in the subsequent year (Grigoryeva and Shatrov, 2022). Delayed life-history effects are generally known to impact dynamics across various species and life histories, and it is particularly important to incorporate such effects in models that aim to predict responses to external impacts (Beckerman et al., 2002). In the case of ixodid ticks, individuals that overwinter without feeding will have depleted more resources next summer compared to ticks that emerge in the same year. This depletion likely affects their survival and potentially their questing behavior as well. Ticks that feed before the summer threshold have a higher probability of molting in the same year, but a lower survival in winter if they have not molted. Ticks that feed after the threshold have a higher probability of survival overwinter, and a low probability of molting until next year. Similarly, ticks that overwinter as fed have a high probability of molting the next summer, and a low probability of survival towards the end of the year. By including such detailed lasting effects, our model obtains a high flexibility to describe a range of scenarios for tick life histories and responses to environmental change.

Arthropod vectors, such as ixodid ticks, are ectotherms that are expanding their distribution towards higher elevations and latitudes under global warming, both in North America and in Europe (Medlock et al., 2013; Clow et al., 2017). Earlier models of *Ixodes* ssp. dynamics have primarily focused on implementing how individual-level processes of questing and development depend on key environmental drivers such as temperature and saturation deficit (Ogden et al., 2005; Dobson et al., 2011; Li et al., 2016). These models generally assumed a constant host availability and emphasized the direct effects of climate drivers as the primary source of seasonality (Ogden et al., 2005; Dobson et al., 2011). However, it is important to consider that the dynamics of small mammalian hosts in Europe have been altered by both land use and climate change (Cornulier et al., 2013), and there has been a marked increase in the distribution and abundance of large host species such as deer in recent decades (Linnell et al., 2020). Our model allows exploring the additional effects of such changes in the host population on the tick dynamics and seasonal abundance.

Our baseline equilibrium analysis for *I. ricinus* reveals several key findings, including that i) small hosts play a more significant role in regulating tick populations compared to large hosts, unless the density of large hosts is exceptionally low, ii) the transition of larvae feeding is critical for population regulation, and iii) the small host preference of larvae, rather than nymphs, is a crucial determinant of tick population regulation. Across all scenarios considered, the density regulation primarily occurs through larval feeding in specific months of high activity (figs. S12-S14, S27-S29, S42-S44). In the baseline northern scenario, a dip in small host feeding by nymphs in June and July (fig. 3b) arises from a combination of factors: a lower number of questing nymphs compared to the peak (fig. 3a), a low probability of finding a small host due to their relative scarcity in June (fig. 1), and competition with larvae for available small hosts. Removing the seasonality of small hosts has small effects on the seasonality of questing ticks (fig. 3a), but it increases the tick population by enabling more ticks to successfully feed during critical questing months (fig. 3b, figs. S27-S29). As a result, the host capacity becomes filled in more months compared to the seasonal scenario. Thus, the effects of changing host seasonality on tick dynamics are complex, as it not only affects the number of ticks that can feed but also potentially influences density-dependent regulation. The results for the southern scenario (Supporting Information S6) highlight the model’s flexibility in capturing diverse tick and host phenology. Unlike the northern scenario, nymphs and larvae in the southern scenario quest at different times with minimal overlap, ensuring that nymphs never compete with larvae for small hosts.

A critical aspect of population models is the detailed processes by which density dependence is incorporated. In ixodid ticks, there is no clear empirical evidence regarding which stage-specific vital rates are most affected by density dependence. However, the feeding process on a limited number of hosts is an evident starting point for including such effects. In our model, we assumed a fixed number of available feeding spots per host, which varied across months, resulting in the host capacity being filled during critical months at equilibrium. Additionally, we included scramble competition among nymphs and larvae for small hosts, while large hosts had separate capacities for each of the main tick stages. By contrast, Dobson et al. (2011) incorporated density dependence in a different manner, specifically through survival during the post-feeding period. In their model, as the number of successfully feeding ticks increased, the survival during the post-feeding period would decline. This mechanism leads to a more immediate mortality effect of high density during feeding compared to our model, where ticks that are unable to feed remain in the unfed stage for a longer duration (thus indirectly increasing the likelihood of mortality before finding a host). Empirical data on tick load on hosts, as well as other parasites, consistently demonstrate uneven distribution (Lindsø et al., 2023). Our model presents a good starting point for incorporating tick aggregation on hosts.

The relative importance of small and large hosts in regulating tick populations and, consequently, disease hazard has received considerable attention due to contrasting patterns reported (Deblinger et al., 1993; Rand et al., 2004; Ostfeld et al., 2006). Adult ticks rely on large hosts for reproduction, making deer potentially crucial amplifiers of the tick population (Ruiz-Fons and Gilbert, 2010; James et al., 2013). Our exploration of varying levels of large host availability (Fig 4) suggests that once a certain number of large hosts are available, increasing their availability further has no effect on the equilibrium tick population. This non-linear effect of deer density is consistent with empirical findings for both *I. scapularis* in North America (Hofmeester et al., 2017) and *I. ricinus* in Europe (Mysterud et al., 2016). The impact of deer density on tick numbers appears weak or even absent after reaching 5-10 deer per km^2^ or higher (Mysterud et al., 2016). Due to their much lower reproductive rate compared to small mammals, we did not consider seasonality in large hosts in our analyses. Deer typically reproduce only once a year, produce few offspring (red deer: 1; roe deer: 2-3), and have a low maximum annual population growth rate (30-35% in red deer, >50% in roe deer). However, deer migration may cause considerable seasonality in local densities (van Moorter et al., 2021). Given that deer numbers had a limited role in regulating tick populations in our scenarios (Fig. 4), we deemed it beyond the scope of this study to incorporate such effects.

When modeling the transmission of pathogens between vectors and hosts, it is crucial to consider the life history and seasonality of hosts (Valenzuela-Sánchez et al., 2021). The current model serves as a good starting point for incorporating such factors. Currently, our model only includes one small host, which could represent multiple small mammals with similar seasonal variations, typically peaking in the fall (Andreassen et al., 2021). However, the role of birds adds further complexity to the system (Loss et al., 2016). Many bird species are latitudinal migrants, reproducing in the spring and peaking in numbers later in the season. Therefore, while our current model accurately captures the broad seasonal patterns of small hosts, it can easily be extended to include more than one small host to capture differences among them, including finer details in their phenology. While we included seasonality of the small host, the numbers each month were kept constant across years and thus represent only a cartoon of the complex dynamics of small mammals, which can show large annual cycles on top of the seasonal variation (Andreassen et al., 2021). Using a mechanistic agent-based model Li et al. (2016) included seasonality and multi-annual cycles in rodent hosts through trigonometric functions. This was a detailed spatial model aimed to predict Lyme disease risk, and they did not explicitly consider the role of host cycles for tick regulation.

Pathogens transmitted by ticks often have a narrower host range compared to the ticks. Although *I. ricinus* is found on a wide range of mammals and birds (Mysterud et al., 2015), relatively few and common species of vertebrates seem to dominate the transmission dynamics of pathogens in Europe (Hofmeester et al., 2016). When incorporating pathogen dynamics into the model, the inclusion of both competent and incompetent (small) reservoir hosts becomes more important, as their relative abundance can lead to host dilution effects (Keesing and Ostfeld, 2021; Ostfeld and Keesing, 2013). Additionally, the aggregation of ticks on hosts, affecting the potential for co-feeding transmission (Brunner and Ostfeld, 2008), could then be explicitly modeled. Likewise, different deer species may not vary much in their role as host to ticks, but vary more in their role for transmission of different pathogens (Fabri et al., 2021). Thus, our model serves as a good starting point for incorporating these factors and can be adapted for a broader range of disease systems.

We have presented a new matrix model framework for investigating the role of underlying mechanisms governing the abundance and seasonality of questing ticks, which is important for disease hazard. The model can be used to explore a range of questions regarding the ecology and evolution of ixodid ticks, including comparative analyses of different tick life histories and environments. Our findings from the scenarios considered emphasize the need for a detailed understanding of vector and host ecology in order to reliably decipher the underlying mechanisms behind the seasonal numbers of questing ticks. Modeling provides an important tool for assessing the relative importance of different drivers such as land use and climate change, with their different impacts across species and regions (Rogers and Randolph, 2006).

## Author contributions

YV defined the model and did the analyses, with inputs from AM. Both authors contributed to write and revise the manuscript.

## References

Andreassen, H. P., J. Sundell, F. Ecke, S. Halle, M. Haapakoski, H. Henttonen, O. Huitu, J. Jacob, K. Johnsen, E. Koskela, J. J. Luque-Larena, N. Lecomte, H. Leirs, J. Mariën, M. Neby, O. Rätti, T. Sievert, G. R. Singleton, J. van Cann, B. Vanden Broecke, and H. Ylönen, 2021. Population cycles and outbreaks of small rodents: Ten essential questions we still need to solve. Oecologia 195:601–622.

Beckerman, A., T. G. Benton, E. Ranta, V. Kaitala, and P. Lundberg, 2002. Population dynamic consequences of delayed life-history effects. Trends in Ecology & Evolution 17:263–269.

Belozerov, V. N., L. J. Fourie, and D. J. Kok, 2002. Photoperiodic control of developmental diapause in nymphs of prostriate ixodid ticks (Acari: Ixodidae). Experimental & Applied Acarology 28:163–168.

Bienvenu, F. and S. Legendre, 2015. A new approach to the generation time in matrix population models. The American Naturalist 185:834–843.

Bregnard, C., O. Rais, C. Herrmann, O. Kahl, K. Brugger, and M. J. Voordouw, 2021. Beech tree masting explains the inter-annual variation in the fall and spring peaks of *Ixodes ricinus* ticks with different time lags. Parasites & Vectors 14:570.

Brunner, J. L. and R. S. Ostfeld, 2008. Multiple causes of variable tick burdens on small-mammal hosts. Ecology 89:2259–2272.

Caswell, H., 2001. Matrix Population Models: Construction, Analysis and Interpretation. Sinauer Associates Inc., Sunderland, Massachussets, second edition.

Caswell, H., 2009. Stage, age and individual stochasticity in demography. Oikos 118:1763–1782.

Clow, K. M., P. A. Leighton, N. H. Ogden, L. R. Lindsay, P. Michel, D. L. Pearl, and C. M. Jardine, 2017. Northward range expansion of *Ixodes scapularis* evident over a short timescale in Ontario, Canada. PLOS ONE 12:e0189393.

Cornulier, T., N. G. Yoccoz, V. Bretagnolle, J. E. Brommer, A. Butet, F. Ecke, D. A. Elston, E. Framstad, H. Henttonen, B. Hörnfeldt, O. Huitu, C. Imholt, R. A. Ims, J. Jacob, B. Jędrzejewska, A. Millon, S. J. Petty, H. Pietiäinen, E. Tkadlec, K. Zub, and X. Lambin, 2013. Europe-wide dampening of population cycles in keystone herbivores. Science 340:63–66.

Dantas-Torres, F. and D. Otranto, 2013. Seasonal dynamics of *Ixodes ricinus* on ground level and higher vegetation in a preserved wooded area in southern Europe. Veterinary Parasitology 192:253–258.

Deblinger, R. D., M. L. Wilson, D. W. Rimmer, and A. Spielman, 1993. Reduced abundance of immature Ixodes dammini (Acari: Ixodidae) following incremental removal of deer. Journal of Medical Entomology 30:144–150.

Dobson, A. D. M., 2014. History and complexity in tick-host dynamics: Discrepancies between ‘real’ and ‘visible’ tick populations. Parasites & Vectors 7:231.

Dobson, A. D. M., T. J. R. Finnie, and S. E. Randolph, 2011. A modified matrix model to describe the seasonal population ecology of the European tick *Ixodes ricinus*. Journal of Applied Ecology 48:1017–1028.

Estrada-Peña, A. and D. Estrada-Sánchez, 2014. Deconstructing *Ixodes ricinus*: A partial matrix model allowing mapping of tick development, mortality and activity rates. Medical and Veterinary Entomology 28:35–49.

Estrada-Peña, A., H. Sprong, A. Cabezas-Cruz, J. de la Fuente, A. Ramo, and E. C. Coipan, 2016. Nested coevolutionary networks shape the ecological relationships of ticks, hosts, and the Lyme disease bacteria of the *Borrelia burgdorferi* (s.l.) complex. Parasites & Vectors 9:517.

Fabri, N. D., H. Sprong, T. R. Hofmeester, H. Heesterbeek, B. F. Donnars, F. Widemo, F. Ecke, and J. P. G. M. Cromsigt, 2021. Wild ungulate species differ in their contribution to the transmission of *Ixodes ricinus*-borne pathogens. Parasites & Vectors 14:360.

Franke, J., A. Hildebrandt, and W. Dorn, 2013. Exploring gaps in our knowledge on Lyme borreliosis spirochaetes – Updates on complex heterogeneity, ecology, and pathogenicity. Ticks and Tick-borne Diseases 4:11–25.

Gatewood, A. G., K. A. Liebman, G. Vourc’h, J. Bunikis, S. A. Hamer, R. Cortinas, F. Melton, P. Cislo, U. Kitron, J. Tsao, A. G. Barbour, D. Fish, and M. A. Diuk-Wasser, 2009. Climate and tick seasonality are predictors of *Borrelia burgdorferi* genotype distribution. Applied and Environmental Microbiology 75:2476–2483.

Gray, J. S., 1981. The fecundity of *Ixodes ricinus* (L.) (Acarina: Ixodidae) and the mortality of its developmental stages under field conditions. Bulletin of Entomological Research 71:533–542.

Gray, J. S., O. Kahl, R. S. Lane, M. L. Levin, and J. I. Tsao, 2016. Diapause in ticks of the medically important *Ixodes ricinus* species complex. Ticks and Tick-borne Diseases 7:992–1003.

Grigoryeva, L. A. and A. B. Shatrov, 2022. Life cycle of the tick Ixodes ricinus (L.) (Acari: Ixodidae) in the north-west of Russia. Systematic and Applied Acarology 27:538–550.

Hill, C. A., F. C. Kafatos, S. K. Stansfield, and F. H. Collins, 2005. Arthropod-borne diseases: Vector control in the genomics era. Nature Reviews Microbiology 3:262–268.

Hofmeester, T. R., E. C. Coipan, S. E. van Wieren, H. H. T. Prins, W. Takken, and H. Sprong, 2016. Few vertebrate species dominate the *Borrelia burgdorferi* s.l. life cycle. Environmental Research Letters 11:043001.

Hofmeester, T. R., H. Sprong, P. A. Jansen, H. H. T. Prins, and S. E. van Wieren, 2017. Deer presence rather than abundance determines the population density of the sheep tick, *Ixodes ricinus*, in Dutch forests. Parasites & Vectors 10:433.

James, M. C., A. S. Bowman, K. J. Forbes, F. Lewis, J. E. Mcleod, and L. Gilbert, 2013. Environmental determinants of *Ixodes ricinus* ticks and the incidence of *Borrelia burgdorferi* sensu lato, the agent of Lyme borreliosis, in Scotland. Parasitology 140:237–246.

Jones, K. E., N. G. Patel, M. A. Levy, A. Storeygard, D. Balk, J. L. Gittleman, and P. Daszak, 2008. Global trends in emerging infectious diseases. Nature 451:990–993.

Kahl, O. and J. S. Gray, 2023. The biology of *Ixodes ricinus* with emphasis on its ecology. Ticks and Tick-Borne Diseases 14:102114.

Keesing, F. and R. S. Ostfeld, 2021. Dilution effects in disease ecology. Ecology Letters 24:2490–2505.

Li, S., L. Gilbert, P. A. Harrison, and M. D. A. Rounsevell, 2016. Modelling the seasonality of Lyme disease risk and the potential impacts of a warming climate within the heterogeneous landscapes of Scotland. Journal of The Royal Society Interface 13:20160140.

Lindsø, L. K., J. L. Anders, H. Viljugrein, A. Herland, V. M. Stigum, W. R. Easterday, and A. Mysterud, 2023. Individual heterogeneity in ixodid tick infestation and prevalence of *Borrelia burgdorferi* sensu lato in a northern community of small mammalian hosts. Oecologia 203:421–433.

Linnell, J. D. C., B. Cretois, E. B. Nilsen, C. M. Rolandsen, E. J. Solberg, V. Veiberg, P. Kaczensky, B. Van Moorter, M. Panzacchi, G. R. Rauset, and B. Kaltenborn, 2020. The challenges and opportunities of coexisting with wild ungulates in the human-dominated landscapes of Europe’s Anthropocene. Biological Conservation 244:108500.

Loss, S. R., B. H. Noden, G. L. Hamer, and S. A. Hamer, 2016. A quantitative synthesis of the role of birds in carrying ticks and tick-borne pathogens in North America. Oecologia 182:947–959.

Mannelli, A., L. Bertolotti, L. Gern, and J. Gray, 2012. Ecology of *Borrelia burgdorferi* sensu lato in Europe: Transmission dynamics in multi-host systems, influence of molecular processes and effects of climate change. FEMS Microbiology Reviews 36:837–861.

Medlock, J. M., K. M. Hansford, A. Bormane, M. Derdakova, A. Estrada-Peña, J.-C. George, I. Golovljova, T. G. T. Jaenson, J.-K. Jensen, P. M. Jensen, M. Kazimirova, J. A. Oteo, A. Papa, K. Pfister, O. Plantard, S. E. Randolph, A. Rizzoli, M. M. Santos-Silva, H. Sprong, L. Vial, G. Hendrickx, H. Zeller, and W. V. Bortel, 2013. Driving forces for changes in geographical distribution of *Ixodes ricinus* ticks in Europe. Parasites & Vectors 6:1–11.

Mysterud, A., R. Byrkjeland, L. Qviller, and H. Viljugrein, 2015. The generalist tick *Ixodes ricinus* and the specialist tick *Ixodes trianguliceps* on shrews and rodents in a northern forest ecosystem - a role of body size even among small hosts. Parasites & Vectors 8:1–10.

Mysterud, A., W. R. Easterday, V. M. Stigum, A. B. Aas, E. L. Meisingset, and H. Viljugrein, 2016. Contrasting emergence of Lyme disease across ecosystems. Nature Communications 7:11882.

Mysterud, A., I. L. Hatlegjerde, and O. J. Sørensen, 2014. Attachment site selection of life stages of *Ixodes ricinus* ticks on a main large host in Europe, the red deer (*Cervus elaphus*). Parasites & Vectors 7:510.

Nyrhilä, S., J. J. Sormunen, S. Mäkelä, E. Sippola, E. J. Vesterinen, and T. Klemola, 2020. One out of ten: Low sampling efficiency of cloth dragging challenges abundance estimates of questing ticks. Experimental and Applied Acarology 82:571–585.

Ogden, N. H., 2017. Climate change and vector-borne diseases of public health significance. FEMS Microbiology Letters 364:fnx186.

Ogden, N. H., M. Bigras-Poulin, C. J. O’Callaghan, I. K. Barker, L. R. Lindsay, A. Maarouf, K. E. Smoyer-Tomic, D. Waltner-Toews, and D. Charron, 2005. A dynamic population model to investigate effects of climate on geographic range and seasonality of the tick *Ixodes scapularis*. International Journal for Parasitology 35:375–389.

Ogden, N. H. and L. R. Lindsay, 2016. Effects of climate and climate change on vectors and vector-borne diseases: Ticks are different. Trends in Parasitology 32:646–656.

Ostfeld, R. S., C. D. Canham, K. Oggenfuss, R. J. Winchcombe, and F. Keesing, 2006. Climate, deer, rodents, and acorns as determinants of variation in Lyme-disease risk. PLOS Biology 4:e145.

Ostfeld, R. S. and F. Keesing, 2013. Straw men don’t get Lyme disease: Response to Wood and Lafferty. Trends in Ecology & Evolution 28:502–503.

Qviller, L., N. Risnes-Olsen, K. M. Bærum, E. L. Meisingset, L. E. Loe, B. Ytrehus, H. Viljugrein, and A. Mysterud, 2013. Landscape level variation in tick abundance relative to seasonal migration in red deer. PLOS ONE 8:e71299.

R Core Team, 2023. R: A language and environment for statistical computing. Technical report, Vienna, Austria.

Rand, P. W., C. Lubelczyk, M. S. Holman, E. H. Lacombe, and R. P. Smith, 2004. Abundance of *Ixodes scapularis* (Acari: Ixodidae) after the complete removal of deer from an isolated offshore island, endemic for Lyme disease. Journal of Medical Entomology 41:779–784.

Randolph, S. E., 2004. Tick ecology: Processes and patterns behind the epidemiological risk posed by ixodid ticks as vectors. Parasitology 129:S37–S65.

Randolph, S. E., R. M. Green, A. N. Hoodless, and M. F. Peacey, 2002. An empirical quantitative framework for the seasonal population dynamics of the tick Ixodesricinus. International Journal for Parasitology 32:979–989.

Reiter, P., 2001. Climate change and mosquito-borne disease. Environmental Health Perspectives 109:141–161.

Rogers, D. J. and S. E. Randolph, 2006. Climate change and vector-borne diseases. Advances in Parasitology 62:345–381.

Ruiz-Fons, F. and L. Gilbert, 2010. The role of deer as vehicles to move ticks, *Ixodes ricinus*, between contrasting habitats. International Journal for Parasitology 40:1013–1020.

Tkadlec, E. and J. Zejda, 1998. Small rodent population fluctuations: The effects of age structure and seasonality. Evolutionary Ecology 12:191–210.

Valenzuela-Sánchez, A., M. Q. Wilber, S. Canessa, L. D. Bacigalupe, E. Muths, B. R. Schmidt, A. A. Cunningham, A. Ozgul, P. T. Johnson, and H. Cayuela, 2021. Why disease ecology needs life-history theory: A host perspective. Ecology Letters 24:876–890.

van Moorter, B., N. J. Singh, C. M. Rolandsen, E. J. Solberg, H. Dettki, J. Pusenius, J. Månsson, H. Sand, J. M. Milner, O. Roer, A. Tallian, W. Neumann, G. Ericsson, and A. Mysterud, 2021. Seasonal release from competition explains partial migration in European moose. Oikos 130:1548–1561.

Wongnak, P., S. Bord, M. Jacquot, A. Agoulon, F. Beugnet, L. Bournez, N. Cèbe, A. Chevalier, J.-F. Cosson, N. Dambrine, T. Hoch, F. Huard, N. Korboulewsky, I. Lebert, A. Madouasse, A. Mårell, S. Moutailler, O. Plantard, T. Pollet, V. Poux, M. René-Martellet, M. Vayssier-Taussat, H. Verheyden, G. Vourc’h, and K. Chalvet-Monfray, 2022. Meteorological and climatic variables predict the phenology of *Ixodes ricinus* nymph activity in France, accounting for habitat heterogeneity. Scientific Reports 12:7833.

Wu-Chuang, A., A. Hodžić, L. Mateos-Hernández, A. Estrada-Peña, D. Obregon, and A. Cabezas-Cruz, 2021. Current debates and advances in tick microbiome research. Current Research in Parasitology & Vector-Borne Diseases 1:100036.

