## Supplementary material for "Unraveling the seasonal dynamics of ixodid ticks: A flexible matrix population model with delayed life history effects": SupplementaryInformation.html

###### January 08 2024

The htlm version of this document presents selected R code, additional analyses and supplementary figures. The R Markdown file includes all code to reproduce the html document.

### Links to figures and tables

The following hyperlinks can be used to navigate directly to each supplementary table/figure.

- Table S1: Overview of stages

**Host availability:**

- Figure S1: Small host vital rates
- Figure S2: Host availability
- Figure S3: Probability of finding host
- Figure S4 (fig. 2): Combined plot, host properties.

**Baseline northern scenario:**

- Figure S5: Parameters
- Figure S6: Projected August population over time
- Figure S7: Monthly stage size at equilibrium
- Figure S8: Stable stage structure
- Figure S9: Reproductive values
- Figure S10: Questing population per month
- Figure S11: Numbers feeding per month per host type
- Figure S12: Proportion of host capacity used
- Figure S13: Proportion of hosts used by each stage
- Figure S14: Stage-specific host capacity used
- Figure S15: Fed population per month (spring, fall, previous)
- Figure S16: Delayed and direct development, including eggs
- Figure S17: Delayed and direct development, excluding eggs
- Figure S18 (fig. 3): Combined plot, results from baseline

**Vary host parameters:**

- Figure S19 (fig. 4): Varying host levels
- Figure S20 (fig. 5): Varying small host preference

**Constant small host availability:**

- Figure S21: Projected August population over time
- Figure S22: Monthly stage size at equilibrium
- Figure S23: Stable stage structure
- Figure S24: Reproductive values
- Figure S25: Questing population per month
- Figure S26: Numbers feeding per month per host type
- Figure S27: Proportion available host capacity used
- Figure S28: Proportion of hosts used by each stage
- Figure S29: Stage-specific host capacity used
- Figure S30: Fed population per month (spring, fall, previous)
- Figure S31: Delayed and direct development, including eggs
- Figure S32: Delayed and direct development, excluding eggs
- Figure S33: Combined plot, results with constant hosts.

**Southern scenario:**

- Figure S34: Survival and density independent transitions
- Figure S35: Comparison of density independent transitions
- Figure S36: Projected August population over time
- Figure S37: Monthly stage size at equilibrium
- Figure S38: Stable stage structure
- Figure S39: Reproductive values
- Figure S40: Questing population per month
- Figure S41: Numbers feeding per month per host type
- Figure S42: Proportion available host capacity used
- Figure S43: Proportion of hosts used by each stage
- Figure S44: Stage-specific host capacity used
- Figure S45: Fed population per month (spring, fall, previous)
- Figure S46: Delayed and direct development, including eggs
- Figure S47: Delayed and direct development, excluding eggs
- Figure S48: Combined plot, results compared to northern
- Figure S49: Combined plot, results southern scenario

### S1 Setup

```
#-------------------
#Packages
#-------------------
library(tidyverse)
library(viridis)
library(cowplot) 
 

#-------------------
#Define stages
#------------------- 
stages <- c("EGG", "EGP", #Egg stages
            "LU",  "LUP", "LFS", "LFF", "LFP", #Larvae stages
            "NU",  "NUP", "NFS", "NFF", "NFP", #Nymph stages
            "AU",  "AUP", "AFS", "AFF", "AFP") #Adult stages

k <- length(stages)  

#-------------------
#Define functions
#-------------------
logit <- function(x){log(x/(1-x))}
inv.logit <- function(x){1/(1+exp(-x))}

#Lambda, stable structure and reproductive values based on eigen analysis:
uvlambda <- function(MatA){
  ev <- eigen(MatA)
  tev <- eigen(t(MatA))
  lmax <- which.max(Re(ev$values))
  U <- ev$vectors
  V <- tev$vectors
  u <- as.matrix(abs(Re(U[, lmax]))/sum(abs(Re(U[, lmax]))))
  u <- u/(sum(u))
  v <- as.matrix(abs(Re(V[, lmax])))
  v <- v/sum(u*v )
  v <- ifelse(u*v <= 0, 0, v)
  v2 <- v/v[1]
  return(list("lambda"=max(Re(ev$values)),"u"=as.vector(u),"v"=as.vector(v),"v2"=as.vector(v2)))
 }
 
#Net reproductive rate:
R0function  <-  function(MatF, MatU){
     k <- dim(MatF)[1] 
     MatR <- MatF%*%solve(diag(1,k,k)-MatU)
     resR<- uvlambda(MatR)
     resR$lambda
 }
 
#Generation time (mean age of mothers at stable distribution):
GenTime  <-  function(MatA, MatF){
  res <- uvlambda(MatA=MatA)
  lam <- res$lambda
  u <- res$u
  v <- res$v
  lam/(t(v)%*%MatF%*%u)    
}
```

##### Overview of stages in the model

Table S1: Overview of tick stages in the model, and the next possible transition out of each stage in a baseline northern scenario. The spring / fall threshold is set to July, individuals feeding in spring must molt to the next stage before winter while individuals feeding after the threshold will overwinter and molt next year. Unfed individuals may also overwinter. Density and host dependence occurs through the feeding process, defining transitions from unfed to fed stages.

| Number | Stage | Description |
| --- | --- | --- |
| 1 | EGG | Egg laid current year |
| 2 | EGP | Egg laid previous year |
| 3 | LU | Unfed larva hatched current year |
| 4 | LUP | Unfed larva hatched previous year |
| 5 | LFS | Larva fed in spring current year |
| 6 | LFF | Larva fed in fall current year |
| 7 | LFP | Larva fed previous year |
| 8 | NU | Unfed nymph emerged current year |
| 9 | NUP | Unfed nymph emerged previous year |
| 10 | NFS | Nymph fed in spring current year |
| 11 | NFF | Nymph fed in fall current year |
| 12 | NFP | Nymph fed previous year |
| 13 | AU | Unfed adult emerged current year |
| 14 | AUP | Unfed adult emerged previous year |
| 15 | AFS | Adult fed in spring current year |
| 16 | AFF | Adult fed in fall current year |
| 17 | AFP | Adult fed previous year |

### S2 Host availability and feeding

We assume that the feeding of ticks is limited by the availability of small hosts (\(S\_i\)) and large hosts (\(L\_i\)). Adult ticks must feed on a large host, while nymphs and larvae can feed on both large and small hosts. Larvae are assumed to prefer small hosts with a high probability (0.9 in the baseline model), while nymphs are more generalists (small host preference 0.5 in the baseline model).

##### Available hosts per month

We assume that large hosts have small or no seasonal fluctuations and thus are available for a larger part of the year, whereas the small hosts show seasonality representative of northern populations.

To describe the seasonal availability we add a simple model for the monthly dynamics of a generic small mammal host (e.g., similar to bank vole), where survival and fecundity change over months but the population is reset every year to the same starting population in January. The small host model assumes a post-reproductive census where individuals are counted in beginning of each month, right after reproduction. Offspring are assumed to be unavailable for ticks, and we assume that individuals reproduce for the first time at the age of two months.

In the alternative scenario for southern Europe, we assume reproduction starts much earlier in the year and ends in summer when plant growth declines due to heat (Andreassen et al., 2021).

Figure S1: Small host vital rates, used to generate seasonal numbers of available small hosts (fig. S3).

Figure S2: Available small hosts and large hosts per month. The small host availability is seasonal in the baseline model and constant in alternative scenarios in our analysis.

##### Probability of finding a host

The probability that a questing tick finds a host depends on the number of available hosts in the area, and is defined as a logistic function as described in the main text. The maximum probability depends on tick stage and host type.

```
p.find.host.small <- function(hosts, max.prob = 0.8, ks =7e-4, H0 = 1500){
  max.prob * (1+exp(-ks*(hosts-H0)))^(-1)
}


p.find.host.large <- function(hosts, max.prob = 0.8, kL =.005, H0 = 100){
 max.prob * (1+exp(-kL*(hosts-H0)))^(-1)
}
```

Figure S3: The probability that a questing tick finds a small or large host as a function of host availability. The maximum probability depends on the stage within each host type, values shown are for the baseline model. Vertical lines indicate the baseline starting value of small hosts in January (left), and the baseline constant values of large hosts (right).

##### Combined plot

Figure S4: Figure 2 in main text.

### S3 Baseline northern scenario

The baseline model represents a ‘northern’ tick life history with high survival rates, slow development rates and short questing season. This is compared to a ‘southern’ life history below with lower survival, faster development and longer questing season.

#### S3.1 Parameters

##### Survival and transitions

The data frame ‘survival’ contains the monthly survival probabilities for each stage. The data frame ‘develop’ contains the density independent probabilities of hatching (for eggs), questing (for unfed stages), molting (fed stages of larvae and nymphs), and reproducing and dying (adult fed females). These probabilities define all transitions except the transitions from unfed to fed stages, which depend on the feeding process described in S2.

Figure S5: Baseline values of survival and density independent probabilities of hatching (eggs), questing (unfed stages), molting (fed stages of larvae and nymphs) and reproducing / dying (adult fed stages). The vertical grey lines indicate the threshold month for spring and fall feeding (July).

##### Egg number

The probability that a fed adult female will reproduce will also affect survival, since reproduction is fatal. We assume each reproducing female lays 1500 eggs, reflecting a typical average value (Gray 1981, Grigoryeva & Shatrov 2002).

#### S3.2 Projection function

The following R function projects the population over `tmax` months. The initial month is set to January unless otherwise specified. If the projection is set to start in another month, the code for plotting below should be modified accordingly to show the right months on the x-axes.

The function returns the monthly values of the size of each stage, the size of the questing population, the size of the fed stages (feeding on small hosts and on large hosts), the size of the emerging population (molting). It also returns the monthly values of the transition matrix, the survival/transition matrix \(\mathbf{U}\), and the fertility matrix \(\mathbf{F}\).

```
Project.pop <- function(tmax=3000, devframe=develop, survframe=survival, eggs_pf=1500, N0=Init, smallhost.pref=c(.9, .5, 0), needed.spots.per.tick.stage = c(1, 3, 10), maxprob.small = c(.7, .9, 0), maxprob.large=c(.3, .6, .9), small.host.capacity = SHCapacity, number.per.host.large = LHFeedingSpots, SpringFallThreshold = 7, start.month=1, HostFrame=Hosts){
  #=====================
  #ORGANIZE PARAMETERS
  #=====================
  stages <- c("EGG", "EGP", 
            "LU",  "LUP", "LFS", "LFF", "LFP",
            "NU",  "NUP", "NFS", "NFF", "NFP",
            "AU",  "AUP", "AFS", "AFF", "AFP")
  unfed.stages <- c("LU", "LUP", "NU", "NUP", "AU", "AUP")
  k <- length(stages)
  #----------------------
  #FERTILITY (Not density dependent)
  #----------------------
  #Array for F-matrices by month
  FmatArray <- array(data = 0, dim=c(12, k, k)) 
  dimnames(FmatArray)[[2]] <-  dimnames(FmatArray)[[3]] <- stages
  for(i in 1:12){
    FmatArray[i,"EGG","AFS"] <- 0.5*eggs_pf*survframe[i,"AFS"]*devframe[i,"AFS"]
    FmatArray[i,"EGG","AFF"] <- 0.5*eggs_pf*survframe[i,"AFF"]*devframe[i,"AFF"]
    FmatArray[i,"EGG","AFP"] <- 0.5*eggs_pf*survframe[i,"AFP"]*devframe[i,"AFP"]
  }
  #----------------------
  #Organize feeding parameters for 6 unfed stages 
  #----------------------
  #LU, LUP, NU, NUP, AU, AUP
  SH.pref <- c(rep(smallhost.pref[1], 2), rep(smallhost.pref[2], 2), 
               rep(smallhost.pref[3], 2)) #Small host preference
  spt <- c(rep(needed.spots.per.tick.stage[1], 2), rep(needed.spots.per.tick.stage[2], 2), 
           rep(needed.spots.per.tick.stage[3], 2))  #Spots per tick needed on small host 
  maxP.small <- c(rep(maxprob.small[1], 2), rep(maxprob.small[2], 2), 
                  rep(maxprob.small[3], 2))  #Max probability of finding small host
  maxP.large <- c(rep(maxprob.large[1], 2), rep(maxprob.large[2], 2),
                  rep(maxprob.large[3], 2))  #Max probability of finding large host
  sph.large <- c(rep(number.per.host.large[1],2), rep(number.per.host.large[2],2), rep(number.per.host.large[3],2)) #Number of ticks that can feed on each large host per month
  #----------------------
  #INITIALIZE
  #----------------------
  Nmat <- matrix(NA, ncol=tmax+1, nrow=k) #To store population size each month
  rownames(Nmat) <- stages
  Nmat[,1] <- N0 
  QuestMat <- matrix(NA, ncol=tmax, nrow=6) #To store questing population
  TMatArray <- array(NA, c(tmax, k, k)) #Store transition matrices (density dependent)
  UMatArray <- array(NA, c(tmax, k, k)) #Store survival/transition matrices (density dependent)
  dimnames(TMatArray)[[2]] <-  dimnames(TMatArray)[[3]] <- stages
  dimnames(UMatArray)[[2]] <-  dimnames(UMatArray)[[3]] <- stages
  FedMatS  <- matrix(NA, ncol=tmax, nrow=6) #store numbers feeding on small hosts 
  FedMatL  <- matrix(NA, ncol=tmax, nrow=6) #store numbers feeding on large hosts 
  rownames(QuestMat) <- rownames(FedMatL) <- rownames(FedMatS) <- unfed.stages
  month <- start.month-1 
  #=====================
  #PROJECTION LOOP 
  #=====================
  for(i in 1:tmax){#Define month within year
    month <- month+1
    if(month > 12){
      month <- 1
    }
    #----------------------
    #TRANSITIONS 
    #----------------------
    TMat <- diag(1, k, k)  
    colnames(TMat) <- rownames(TMat) <- stages
    #---------------------
    #Feeding transitions (Density dependent)
    #---------------------
    #Number of unfed ticks per stage
    Unfed <- Nmat[unfed.stages, i]
    #Questing probabilities per stage
    qprob <- devframe[month, unfed.stages]
    #~~~~~~~~~~
    #SMALL HOSTS
    #~~~~~~~~~~
    #Host availability
    HS <- HostFrame$SmallHosts[month] 
    #Number of ticks questing and finding a small host:
    nfind.small <- Unfed * qprob * SH.pref * p.find.host.small(HS, max.prob = maxP.small)
    #Available feeding spots
    feeding.spots.small <- small.host.capacity * HS 
    #Requested feeding spots 
    nreq.small <- nfind.small * spt
    #Number of fed ticks on small
    smallhostfed <- {#Sucessful feeding
      if(sum(nreq.small) > feeding.spots.small){#if not enough spots
        (nreq.small / sum(nreq.small)) * (feeding.spots.small / spt)
        } else {#if enough spots
        nreq.small / spt
        } 
      }
    #~~~~~~~~~~
    #LARGE HOSTS
    #~~~~~~~~~~
    #Host availability 
    HL <- HostFrame$LargeHosts[month] 
    #Number of ticks questing and finding a large host:
    nfind.large <- Unfed * qprob * (1 - SH.pref) * p.find.host.large(HL, max.prob = maxP.large)
    #Available feeding spots per stage
    feeding.spots.large <- sph.large*HL
    largehostfed <- smallhostfed #initiate vector
    for(j in 1:length(nfind.large)){
    if(nfind.large[j] > (feeding.spots.large)[j]){#not enough spots
      largehostfed[j] <- (feeding.spots.large/sum(sph.large))[j]
      }
    if(nfind.large[j] <= (feeding.spots.large)[j]){#enough spots
      largehostfed[j] <- nfind.large[j]
      }
    }
    #Total numbers of fed ticks
    fedticks <- largehostfed + smallhostfed #(before survival)
    FedMatS[, i] <- t(smallhostfed) #store fed on small (before survival)
    FedMatL[, i] <- t(largehostfed) #store fed on large (before survival)
    QuestMat[, i] <- t(Unfed*qprob) #store questing (before survival)
    tprob <- ifelse(Unfed > 0, fedticks/Unfed, 0)
  #----------------------------------
  #Hatching / molting probabilities
  #----------------------------------
  TMat["LU", "EGG"]  <-  devframe[month, "EGG"]   
  TMat["EGG", "EGG"] <-  1 - devframe[month, "EGG"]  
  TMat["LU", "EGP"]  <-  devframe[month, "EGP"] 
  TMat["EGP", "EGP"] <-  1 - devframe[month, "EGP"]  
  TMat["NU", "LFP"]  <-  devframe[month, "LFP"]  
  TMat["LFP", "LFP"] <-  1 - devframe[month, "LFP"] 
  TMat["NU", "LFS"]  <-  devframe[month, "LFS"]  
  TMat["LFS", "LFS"] <-  1 - devframe[month, "LFS"] 
  TMat["NU", "LFF"]  <-  devframe[month, "LFF"]  
  TMat["LFF", "LFF"] <-  1 - devframe[month, "LFF"] 
  TMat["AU", "NFP"]  <-  devframe[month, "NFP"]   
  TMat["NFP", "NFP"] <-  1 - devframe[month, "NFP"]  
  TMat["AU", "NFS"]  <-  devframe[month, "NFS"]   
  TMat["NFS", "NFS"] <-  1 - devframe[month, "NFS"] 
  TMat["AU", "NFF"]  <-  devframe[month, "NFF"] 
  TMat["NFF", "NFF"] <-  1 - devframe[month, "NFF"] 
  TMat["AFS", "AFS"] <-  1 - devframe[month, "AFS"]#Not reproducing
  TMat["AFF", "AFF"] <-  1 - devframe[month, "AFF"]#Not reproducing
  TMat["AFP", "AFP"] <-  1 - devframe[month, "AFP"]#Not reproducing
  if(month < SpringFallThreshold){#FEEDING
    TMat["LFS", "LU"]  <-  tprob$LU 
    TMat["LU", "LU"]   <-  1 - tprob$LU 
    TMat["LFS", "LUP"] <-  tprob$LUP
    TMat["LUP", "LUP"] <-  1 - tprob$LUP 
    TMat["NFS", "NU"]  <-  tprob$NU 
    TMat["NU", "NU"]   <-  1 -  tprob$NU
    TMat["NFS", "NUP"] <-  tprob$NUP
    TMat["NUP", "NUP"] <-  1 -  tprob$NUP
    TMat["AFS", "AU"]  <-  tprob$AU    
    TMat["AU", "AU"]   <-  1 - tprob$AU  
    TMat["AFS", "AUP"] <-  tprob$AUP    
    TMat["AUP", "AUP"] <-  1 - tprob$AUP  
    }
  if(month >= SpringFallThreshold){
    TMat["LFF", "LU"]  <-  tprob$LU 
    TMat["LU", "LU"]   <-  1 - tprob$LU 
    TMat["LFF", "LUP"] <-  tprob$LUP  
    TMat["LUP", "LUP"] <-  1 - tprob$LUP  
    TMat["NFF", "NU"]  <-  tprob$NU   
    TMat["NU", "NU"]   <-  1 - tprob$NU  
    TMat["NFF", "NUP"] <-  tprob$NUP  
    TMat["NUP", "NUP"] <-  1 - tprob$NUP  
    TMat["AFF", "AU"]  <-  tprob$AU 
    TMat["AU", "AU"]   <-  1 - tprob$AUP
    TMat["AFF", "AUP"] <-  tprob$AUP
    TMat["AUP", "AUP"] <-  1 - tprob$AUP 
    }
if(month==12){#Transitions Dec/Jan including shifts to "previous"
  TMat <- diag(0, k, k) 
  colnames(TMat) <- rownames(TMat) <- stages
  TMat["EGP", "EGG"] <- 1 -  devframe[month, "EGG"]    
  TMat["LUP", "EGG"] <- devframe[month,"EGG"]
  TMat["EGP", "EGP"] <- 1 - devframe[month,"EGP"]
  TMat["LUP", "EGP"] <- devframe[month,"EGP"]
  TMat["LUP", "LU"] <-  1 - tprob$LU
  TMat["LFP", "LU"] <-  tprob$LU
  TMat["LUP", "LUP"] <- 1 - tprob$LUP
  TMat["LFP", "LUP"] <- tprob$LUP
  TMat["LFP", "LFF"] <-  1 - devframe[month,"LFF"]
  TMat["NUP", "LFF"] <-  devframe[month,"LFF"]
  TMat["LFP", "LFS"] <-  1 - devframe[month,"LFS"]
  TMat["NUP", "LFS"] <-  devframe[month,"LFS"]
  TMat["LFP", "LFP"] <- 1 - devframe[month,"LFP"]
  TMat["NUP", "LFP"] <- devframe[month,"LFP"]
  TMat["NUP", "NU"] <-  1 - tprob$NU
  TMat["NFP", "NU"] <-  tprob$NU
  TMat["NUP", "NUP"] <- 1-tprob$NUP
  TMat["NFP", "NUP"] <- tprob$NUP
  TMat["NFP", "NFF"] <-  1 - devframe[month,"NFF"]
  TMat["AUP", "NFF"] <-  devframe[month,"NFF"]
  TMat["NFP", "NFS"] <-  1 - devframe[month,"NFS"]
  TMat["AUP", "NFS"] <-  devframe[month,"NFS"]
  TMat["NFP", "NFP"] <- 1-devframe[month,"NFP"]
  TMat["AUP", "NFP"] <- devframe[month,"NFP"]
  TMat["AUP", "AU"] <-  1 - tprob$AU
  TMat["AFP", "AU"] <-  tprob$AU
  TMat["AUP", "AUP"] <- 1-tprob$AUP
  TMat["AFP", "AUP"] <- tprob$AUP 
  TMat["AFP", "AFS"] <-  1 - devframe[month, "AFS"]
  TMat["AFP", "AFF"] <-  1 - devframe[month, "AFF"]
  TMat[ "AFP", "AFP"] <-  1 - devframe[month, "AFP"]
  }
  TMatArray[i,,] <- TMat #Store current transition matrix
  #PROJECTION MATRIX
    UMat <- TMat * as.numeric(t(matrix(survframe[month, 3:(k+2)], ncol=k, nrow=k)))
    UMatArray[i,,] <- UMat #Store current survival/transition matrix
    FMat <- FmatArray[month,,] 
    AMat <- UMat + FMat 
    Nmat[,i+1] <- Nmat[,i]%*%t(AMat) #Project population to next month
  }#end loop over months
  namemonths <- c(month.abb[start.month:12],rep(month.abb,tmax+1))[1:(tmax+1)]
  TotalPop <- data.frame(Nmat)
  Questing <- data.frame(QuestMat)
  FedOnSmall <- data.frame(FedMatS)
  FedOnLarge <- data.frame(FedMatL)
  names(TotalPop) <- namemonths   
  names(Questing) <-   names(FedOnSmall) <-   names(FedOnLarge) <-   namemonths[1:tmax]
  dimnames(TMatArray)[[1]] <- namemonths[1:tmax]
  dimnames(UMatArray)[[1]] <- namemonths[1:tmax]
  return(list("Pop" = TotalPop, 
              "QuestingPop" = Questing, 
              "FedOnSmallHost" = FedOnSmall, 
              "FedOnLargeHost" = FedOnLarge,
              "FMatArray" = FmatArray,
              "UMats" = UMatArray,
              "TMats" = TMatArray))
  }
```

#### S3.3 Run the baseline model

```
SHLevel <- 10000 #Starting value in January, small host adults
LHLevel <- 500 #Large host level
SHCapacity <- 500 #Capacity of small hosts to feed  larvae and nymphs per month
LHFeedingSpots <- c(5000, 5000, 5000) #Number of larvae, nymphs and adults that can feed on each large host per month

Hosts <- data.frame(Month = month.abb)
Hosts$LargeHosts <- rep(1,12)*LHLevel
Hosts$SmallHosts <- popsize.SH(N0=SHLevel, VR=vitalrates.SH)$Adults
Hosts$Month <- factor(Hosts$Month,levels=month.abb)

run <- Project.pop(tmax=3000, devframe=develop, survframe=survival, eggs_pf=1500, N0=Init, smallhost.pref=c(.9, .5, 0), needed.spots.per.tick.stage = c(1, 3, 10), maxprob.small = c(.7, .9, 0), maxprob.large=c(.3, .6, .9), small.host.capacity = SHCapacity, number.per.host.large = LHFeedingSpots, SpringFallThreshold = 7, start.month=1, HostFrame=Hosts)
```

##### Projected August population

August was arbitrarily selected, this plot shows how the projection approaches a stable equilibrium. With the initial values used this takes around 50 years.

Figure S6: August population for year 1 to 200 in the projection run with baseline values.

##### Equilibrium population

Extract the monthly stage size after 198, 199 and 200 years (to verify these are the same).

Figure S7: Size of each stage against month at equilibrium in the baseline model. The vertical grey lines indicate the threshold month for spring and fall feeding (July).

##### Life history outputs at equilibrium

The population growth rate is 1, net reproductive rate is 1, generation time is 3.72 years.

Figure S8: Stable structure at equilibrium, calculated from annual projection matrices starting at each month, for the baseline model.

Figure S9: Reproductive values at equilibrium, calculated from annual projection matrices starting at each month, for the baseline model.

In general, most of the tick population is found as eggs or unfed larvae. The reproductive value is highest for fed adults, which are close to reproduction. Adults fed in spring are in the model assumed to die before winter if they do not reproduce, so have lower reproductive values that adults fed in fall or in the previous year. Unfed adults also have quite high reproductive value.

##### Questing population

Figure S10: Questing population of each stage per month at equilibrium, in the baseline model. The vertical grey lines indicate the threshold month for spring and fall feeding (July).

##### Feeding individuals per month

Figure S11: Numbers feeding on each host type (small or large) per month at equilibrium, in the baseline model. The vertical grey lines indicate the threshold month for spring and fall feeding (July).

##### Host use

What proportion of the total available feeding spots are used per month, by all ticks irrespective of stage?

Figure S12: The proportion of available host capacity used per month, for small and large hosts. Calculated as number of used feeding spots divided by number of available feeding spots per host type per month.

###### Host use per stage

What proportion of the total available feeding spots are used per tick stage per month at equilibrium?

Figure S13: The proportion of available host capacity used per stage, for small and large hosts. The sum of the stage-specific proportions equals the total proportion of the host capacity used.

What proportion of available feeding spots to each tick stage are used by that stage per month?

Figure S14: The proportion of available host capacity for each stage that is used per stage, for small and large hosts. Calculated for each host type as the number of used feeding spots per stage divided by number of available feeding spots per stage per month.

##### Fed population

How do the fed stages (LFS, LFF, LFP, NFS, NFF, NFP, AFS, AFF, AFP) change throughout the year at equilibrium?

Figure S15: Size of the spring and fall fed population (larvae, nymphs and adults) in years 198 to 200.

##### Direct and delayed development

In the new generation of unfed individuals that appear each year (stages EGG, LU, NU, and AU), how many are produced from individuals that fed last fall or eggs that were laid last fall (previous, representing delayed development) versus from individuals feeding or eggs laid in the current year (representing direct development)?

Figure S16: Eggs laid from adults fed current year or previous year, and newly emerged individuals developed from individuals feeding that fed in the current or previous year. The vertical grey lines indicate the threshold month for spring and fall feeding (July).

Figure S17: Newly emerged individuals developed from individuals feeding that fed in the current or previous year. The vertical grey lines indicate the threshold month for spring and fall feeding (July).

##### Combined plot

Figure S18: Results from the baseline model, figure 3 in main text. A. Questing population (current/previous), B. Numbers feeding per month (on small/large host), C. Fed population from spring, fall, and from previous year, and D. Emerging unfed individuals per month (direct / delayed development). The vertical grey lines indicate the threshold month for spring and fall feeding (July).

### S4 Vary host parameters

#### S4.1 Vary host levels

In these analyses we first vary the small host size in January (small host level), or the large host level (which is constant across months).

Figure S19: Questing population with varying small host level (A) and varying large host level (B), all other parameters are the same as in the baseline model

#### S4.2 Vary small host preference

In these analyses we vary the small host preference parameter of larvae and nymphs, one by one. In the baseline model the small host preference is 0.9 in larvae and 0.5 in nymphs.

Figure S20: Questing population with varying small host preference in nymphs (A) and in larvae (B). All other parameters are the same as in the baseline model.

### S5 Constant small host

Here we use the same parameters as in the baseline scenario, except that the small host availability is constant across months (equal to the mean availability of the baseline scenario). Some results are plotted with the results from the baseline scenario (S3) as comparison.

```
HostsConstant <- data.frame(Month = month.abb)
HostsConstant$LargeHosts <- rep(1,12)*LHLevel
HostsConstant$SmallHosts <- popsize.SH(N0=SHLevel*.9769*2, VR=vitalrates.SH.Const, Constant=TRUE)$Adults
HostsConstant$Month <- factor(HostsConstant$Month,levels=month.abb)

run2 <- Project.pop(tmax=3000, devframe=develop, survframe=survival, eggs_pf=1500, N0=Init, smallhost.pref=c(.9, .5, 0), needed.spots.per.tick.stage = c(1, 3, 10), maxprob.small = c(.7, .9, 0), maxprob.large=c(.3, .6, .9), small.host.capacity = SHCapacity, number.per.host.large = LHFeedingSpots, SpringFallThreshold = 7, start.month=1, HostFrame=HostsConstant)
```

##### Projected August population

Figure S21: Equilibrium population size, in the model with constant small host availability compared to the baseline model with seasonal small host availability.

##### Equilibrium population

Figure S22: Size of each stage against month at equilibrium, in the model with constant small host availability compared to the baseline model with seasonal small host availability. The vertical grey lines indicate the threshold month for spring and fall feeding (July).

##### Life history outputs

The population growth rate is 1, net reproductive rate is 1, and the generation time is 3.61 years (compared to 3.72 with seasonal host).

Figure S23: Stable structure at equilibrium, calculated from annual projection matrices starting at each month, for the baseline model with constant small host.

Figure S24: Reproductive values at equilibrium, calculated from annual projection matrices starting at each month, for the baseline model with constant small host.

##### Questing population

Figure S25: Questing population of each stage per month at equilibrium, in the model with constant small host availability compared to the baseline model with seasonal small host availability. The vertical grey lines indicate the threshold month for spring and fall feeding (July).

##### Feeding individuals per month

Figure S26: Size of the fed population (larvae, nymphs and adults) on each host type (small or large) per month at equilibrium, in the model with constant small host availability compared to the baseline model with seasonal small host availability. The vertical grey line indicates the threshold month for spring and fall feeding (July).

##### Host use

Figure S27: The proportion of available host capacity used per month, for small and large hosts, compared to the scenario with seasonal small host. Calculated as number of used feeding spots divided by number of available feeding spots per host type per month.

###### Host use per stage

Figure S28: The proportion of available host capacity used per stage, for small and large hosts, compared to the scenario with seasonal small host. The sum of the stage-specific proportions equals the total proportion of the host capacity used.

Figure S29: The proportion of available host capacity for each stage that is used per stage, for small and large hosts, compared to the scenario with seasonal small host. Calculated for each host type as the number of used feeding spots per stage divided by number of available feeding spots per stage per month.

##### Fed population

Figure S30: Size of the spring and fall fed population (larvae, nymphs and adults) at equilibrium, in the model with constant small host availability compared to the baseline model with seasonal small host availability. The vertical grey line indicates the threshold month for spring and fall feeding (July).

##### Direct and delayed development

Figure S31: Eggs laid from adults fed current year or previous year, and newly emerged individuals developed from individuals feeding that fed in the current or previous year, at equilibrium in the model with constant small host availability. The vertical grey line indicates the threshold month for spring and fall feeding (July).

Figure S32: Newly emerged individuals developed from individuals feeding that fed in the current or previous year at equilibrium, in the model with constant small host availability compared to the baseline model with seasonal small host availability. The vertical grey line indicates the threshold month for spring and fall feeding (July).

##### Combined plot

Figure S33: Equilibrium results in the model with constant small host availability compared to the baseline model with seasonal small host availability. A. Questing population (current/previous), B. Numbers feeding per month (on small/large host), C. Fed population from spring, fall, and from previous year, and D. Emerging unfed individuals per month (direct / delayed development). The vertical grey lines indicate the threshold month for spring and fall feeding (July).

### S6 Southern scenario

In this scenario we consider a southern life history assumed to represent the southern part of the distribution range of *Ixodes ricinus* in Europe. The parameters are chosen to fit the questing phenology reported by Dantas-Torres et al. 2013, and we assume lower survival than in the baseline northern scenario, due to increased risk of desiccation as described by Estrada-Peña & Estrada-Sanchez (2014). We do not know the probabilities for molting and reproduction, but we assume that development progresses overall faster than in northern poplations due to higher mean temperature in all months.

For this model we assume that all larvae quest in the first year when they are hatched (no overwintering of unfed larvae). The spring/fall treshold is set to August, but does not affect survival of spring fed individuals (as in the Northern model) and the likelihood of molting is also set to be the same. In other words, adjusting the spring / fall threshold has no influence on this population.

#### S6.1 Parameters

##### Survival and transitions

Figure S34: Baseline values of survival and density independent transition probabilities in the different stages, for the Southern life history. All individuals die after reproducing so in reproductive stages (‘AFS’, ‘AFF’, ‘AFP’) the transition probability corresponds to the probability of reproducing and dying. Seasonal density independent baseline probabilities of hatching (eggs), questing (unfed stages), molting (fed stages of larvae and nymphs) and reproducing / dying (adult fed stages). Feeding is a density dependent process depending on host availability in addition to questing probability (specified below).

Figure S35: Comparison of the density independent transition probabilities in the baseline Northern and the alternative Southern life history.

##### Egg number per female

We assume the same fecundity, 1500 eggs per female, as in the baseline scenario.

#### S6.2 Run the model

```
HostsSouthern <- data.frame(Month = month.abb)
HostsSouthern$LargeHosts <- rep(1,12)*LHLevel
HostsSouthern$SmallHosts <-  popsize.SH(N0=SHLevel*.55,VR=vitalrates.SH.Southern)$Adults
HostsSouthern$Month <- factor(Hosts$Month,levels=month.abb)

run.southern <- Project.pop(tmax=3000, devframe=develop.southern, survframe=survival.southern, eggs_pf=1500, N0=Init, smallhost.pref=c(.9, .5, 0), needed.spots.per.tick.stage = c(1, 3, 10), maxprob.small = c(.7, .9, 0), maxprob.large=c(.3, .6, .9), small.host.capacity = SHCapacity, number.per.host.large = LHFeedingSpots, SpringFallThreshold = 8, start.month=1, HostFrame = HostsSouthern)
```

##### Projected August population

Figure S36: Equilibrium population size in the Southern model with constant small host, compared to the baseline Northern model including small host seasonality.

##### Equilibrium population

Figure S37: Equilibrium population size in the Southern model with constant small host, compared to the baseline Northern model with seasonal small host. The vertical grey lines indicate the threshold month for spring and fall feeding (July for Northern, August for Southern).

##### Life history outputs

The population growth rate is 1, net reproductive rate is 1, and generation time is 2.23 years (compared to 3.72 in the Northern baseline scenario).

Figure S38: Stable structure at equilibrium, calculated from annual projection matrices starting at each month, for the southern scenario.

Figure S39: Reproductive values at equilibrium, calculated from annual projection matrices starting at each month, for the southern scenario.

##### Questing population

Figure S40: Questing population of each stage per month at equilibrium, comparing the Northern baseline model with small host seasonality to the alternative Southern scenario with constant small host. The vertical grey lines indicate the threshold month for spring and fall feeding (July for Northern, August for Southern).

##### Feeding population per month

Figure S41: Size of the fed population (larvae, nymphs and adults) on each host type (small or large) per month at equilibrium, comparing the Northern baseline model with small host seasonality to the alternative Southern scenario with constant small host. The vertical grey lines indicate the threshold month for spring and fall feeding (July for Northern, August for Southern).

##### Host use

Figure S42: The proportion of available host capacity used per month, for small and large hosts, compared to the baseline northern scenario. Calculated as number of used feeding spots divided by number of available feeding spots per host type per month.

###### Host use per stage

Figure S43: The proportion of available host capacity used per stage, for small and large hosts, compared to the baseline northern scenario. The sum of the stage-specific proportions equals the total proportion of the host capacity used.

Figure S44: The proportion of available host capacity for each stage that is used per stage, for small and large hosts, compared to the baseline northern scenario. Calculated for each host type as the number of used feeding spots per stage divided by number of available feeding spots per stage per month.

##### Fed population

Figure S45: Size of the spring and fall fed population (larvae, nymphs and adults) at equilibrium, comparing the Northern baseline model with small host seasonality to the alternative Southern scenario with constant small host. The vertical grey lines indicate the threshold month for spring and fall feeding (July for Northern, August for Southern).

##### Direct and delayed development

Figure S46: Eggs laid from adults fed current year or previous year, and newly emerged individuals developed from individuals feeding that fed in the current or previous year at equilibrium in the southern scenario, compared to the northern scenario. The vertical grey lines indicate the threshold month for spring and fall feeding (July for Northern, August for Southern).

Figure S47: Newly emerged individuals developed from individuals feeding that fed in the current or previous year at equilibrium, comparing the Northern baseline model with small host seasonality to the alternative Southern scenario with constant small host. The vertical grey lines indicate the threshold month for spring and fall feeding (July for Northern, August for Southern).

##### Combined plot

Figure S48: Equilibrium results for the Northern baseline model with small host seasonality compared to the alternative Southern scenario with constant small host. A. Questing population (current/previous), B. Numbers feeding per month (on small/large host), C. Fed population from spring, fall, and from previous year, and D. Emerging unfed individuals per month (direct / delayed development). The vertical grey lines indicate the threshold month for spring and fall feeding (July for Northern, August for Southern).

Figure S49: Equilibrium results for the southern scenario alone. A. Questing population (current/previous), B. Numbers feeding per month (on small/large host), C. Fed population from spring, fall, and from previous year, and D. Emerging unfed individuals per month (direct / delayed development). The vertical grey lines indicate the threshold month for spring and fall feeding (August).

### References

Andreassen, H.P., Sundell, J., Ecke, F., Halle, S., Haapakoski, M., Henttonen, H., Huitu, O., Jacob, J., Johnsen, K., Koskela, E., Luque-Larena, J.J., Lecomte, N., Leirs, H., Mariën, J., Neby, M., Rätti, O., Sievert, T., Singleton, G.R., van Cann, J., Vanden Broecke, B., Ylönen, H., 2021. Population cycles and outbreaks of small rodents: Ten essential questions we still need to solve. Oecologia 195, 601–622. https://doi.org/10.1007/s00442-020-04810-w

Dantas-Torres, F., Otranto, D., 2013. Seasonal dynamics of *Ixodes ricinus* on ground level and higher vegetation in a preserved wooded area in southern Europe. Vet. Parasitol. 192, 253–258. https://doi.org/10.1016/j.vetpar.2012.09.034

Estrada-Peña, A., Estrada-Sánchez, D., 2014. Deconstructing *Ixodes ricinus*: a partial matrix model allowing mapping of tick development, mortality and activity rates. Med. Vet. Entomol. 28, 35–49. https://doi.org/10.1111/mve.12009

Gray, J.S., 1981. The fecundity of *Ixodes ricinus* (L.) (Acarina: Ixodidae) and the mortality of its developmental stages under field conditions. Bull. Entomol. Res. 71, 533–542. https://doi.org/10.1017/S0007485300008543

Grigoryeva, L.A., Shatrov, A.B., 2022. Life cycle of the tick *Ixodes ricinus* (L.) (Acari: Ixodidae) in the north-west of Russia. Syst. Appl. Acarol. 27, 538–550. https://doi.org/10.11158/saa.27.3.11
